# Nuclear pores as conduits for fluid flow during osmotic stress

**DOI:** 10.1101/2024.01.17.575985

**Authors:** Patrick C. Hoffmann, Hyuntae Kim, Agnieszka Obarska-Kosinska, Jan Philipp Kreysing, Eli Andino-Frydman, Sergio Cruz-Leon, Lenka Cernikova, Jan Kosinski, Beata Turoňová, Gerhard Hummer, Martin Beck

**Affiliations:** Department of Molecular Sociology, Max Planck Institute of Biophysics, Max-von-Laue-Straße 3, 60438 Frankfurt am Main, Germany; Department of Theoretical Biophysics, Max Planck Institute of Biophysics, Max-von-Laue-Straße 3, 60438 Frankfurt am Main, Germany; IMPRS on Cellular Biophysics, Max-von-Laue-Straße 3, 60438 Frankfurt am Main, Germany; European Molecular Biology Laboratory Hamburg, 22607 Hamburg, Germany; Institute of Biophysics, Goethe University Frankfurt, 60438 Frankfurt am Main, Germany; Institute of Biochemistry, Goethe University Frankfurt, 60438 Frankfurt am Main, Germany

## Abstract

Changing environmental conditions necessitate an immediate cellular adaptation to ensure survival. *Dictyostelium discoideum*, a bacteriovore slime mold present in the soil of most terrestrial ecosystems, is known for its ability to tolerate drastic changes in osmolarity. How the cells cope with the resulting mechanical stress remains understudied. Here we show that *D. discoideum* has extraordinarily elaborate and resilient nuclear pores that serve as conduits for massive fluid exchange between cytosol and nucleus. We capitalize on the unique properties of *D. discoideum* cells to quantify flow across the nuclear envelope that is necessitated by changing nuclear size in response to osmotic stress. Based on mathematical concepts adapted from hydrodynamics, we conceptualize this phenomenon as porous flow across nuclear pores. This type of fluid flow is distinct from the canonically characterized modes of nucleocytoplasmic transport, i.e. passive diffusion and active nuclear transport, because of its dependence on pressure. Our insights are relevant in any biological condition that necessitates rapid nuclear size changes, which includes metastasizing cancer cells squeezing through constrictions, migrating cells and differentiating tissues.

## Introduction

Numerous organisms adapt efficiently to rapid environmental changes. Understanding the mechanisms underlying these biological adaptation processes may become crucial to devise strategies to cope with ongoing changes in the global environment. *Dictyostelium discoideum* is a soil-dwelling amoebae that evolved to tolerate drastic alterations in the environmental conditions as a result of seasonal changes. Slime molds require water films for survival at different stages of their life cycle and development ^1,2^. Therefore, they have successfully established means to endure fluctuations in humidity and solute availability. Generally, cells without the additional support of a cell wall react to osmotic gradients with swelling or shrinkage through influx or efflux of water. In response, cells stabilize and regulate their volume by transporting ions and organic osmolytes across their membranes ^3,4^. In addition, some protists including *D. discoideum* have a specialized organelle, the contractile vacuole. The contractile vacuole helps to maintain the osmotic homeostasis of the cell and to cope with conditions of extreme osmolarity. In hypotonic environments, the vacuole removes water from the cytosol and is periodically expelled from the cell when fluid-filled ^5,6^. *D. discoideum* is known to adjust its cellular volume upon osmotic changes on remarkable spatiotemporal scales, which coincide with alterations to cytoskeleton and activation of signaling pathways ^7–11^.

The induced volume changes affect the concentration of all intra-cellular constituents including macromolecules ^12^. Hyperosmotic stress is known to promote molecular crowding, influencing diffusion of molecules, viscosity and molecular condensation ^13,14^. Organelles such as the nucleus respond to the altered cellular volume. Under normal conditions nuclear size is coupled to cell size ^15^. The initial hypothesis that the ratio is mainly linked to DNA content and chromatin compaction has been extended, as it was recently reported that the balance of colloid osmotic pressure of macromolecules localized in the nucleoplasm and cytoplasm determines the volume ratio of the two compartments ^16,17^. The nucleus as a membrane-separated compartment is covered with nuclear pores complexes (NPCs) and as such displays different membrane properties compared to other organellar membranes ^18,19^. For this reason, alterations to cell volume and macromolecular concentration that impact on nuclear shape and volume, should induce compensational fluid flow into or out of the nucleus. How the magnitude and timing of such nuclear size adjustments is determined, and if NPCs contribute to the resulting flow and the maintenance of nuclear homeostasis is currently not clear.

NPCs are large membrane protein complexes embedded in the nuclear membranes and consist of over 1000 copies of ∼30 individual nucleoporin proteins (Nups) (reviewed in ^20–23^). The central channel of the NPC functions as a permeability barrier between cyto- and nucleoplasm and facilitates two well-characterized types of nucleocytoplasmic exchange: active transport against concentration gradients and passive diffusion ^24–26^. The FG-repeat containing Nups (FG-Nups) constitute a permeability barrier that allows passive diffusion of small molecules, while limiting the passage of macromolecules larger than 30 to 40 kDa ^27–29^. Larger cargo molecules above the barrier threshold for macromolecules rely on interactions with nuclear transport receptors for active transport through the central channel. Recent experiments however suggest that the permeability barrier and its threshold function may be more dynamic ^30^. The exact organization of FG-Nups within the permeability barrier of the central channel remains subject to investigation ^24,31^, however, the passive diffusion limit of cargo suggests a characteristic pore size radius of around 2.6 nm in the FG-Nup mesh ^27^.

For several species, including human and yeast, the architecture of the structured NPC scaffold and its components has been studied extensively by X-ray crystallography, electron cryo-tomography (cryo-ET) and integrative modeling ^32,33^. Meanwhile, an almost complete picture of the structured NPC scaffold in these systems has been obtained ^34–38^. The NPC scaffold has an eight-fold rotational symmetry and can be subdivided into three major parts, the cytosolic ring (CR), the inner ring (IR) and the nuclear ring (NR) ^39^. NPCs are composed of different subcomplexes, of which the Y-complex ^40,41^ is the most prominent and structurally best characterized constituent. Y-complexes form concentric rings in a head-to-tail arrangement for both NR and CR ^39^. The eight spokes of the IR are anchored in the NE membrane and provide a binding platform for central channel FG-repeat containing Nups ^42–44^. At last, a single Nup termed Nup210 forms a ring-shaped domain in the nuclear envelope lumen ^45,46^.

Recent studies in cells found that NPCs are conformationally highly dynamic. In unperturbed exponentially growing cells, they reside in a so-called dilated state ^47–51^. In response to environmental cues such as energy depletion and hyperosmotic stress, NPCs constrict into a conformational ground state ^47^. Osmotic gradients have been linked to nuclear membrane tension, which is coupled to NE thickness and the NPC diameter ^47^. It has been proposed that the forces imposed by increased membrane tension stretch the NPC scaffold thus leading to NPC dilation, while in the absence of tension NPCs are constricted ^47^. Besides alterations in nucleocytoplasmic transport ^47,52^, the exact functional relevance of NPC diameter adaptation remains understudied.

Here, we present the *in situ* structure of the *D. discoideum* NPC scaffold and its changes upon osmotic stress. We capitalize on the unique properties of *D. discoideum* cells to quantify flow across the nuclear envelope that is necessitated by changes in nuclear size. We propose that it is best conceptualized as porous flow across nuclear pores.

## Results

### Cellular and nuclear volume adaptations occur within seconds to minutes in response to osmotic stress

In eukaryotes, nuclear and cell size are coupled under normal growth conditions ^15^. Although osmotic stress has immediate and drastic effects on the overall cell size of *D. discoideum* ^8,11^, it yet remains unclear on what timescale such acute cell size changes are translated to the nucleus. To experimentally address this, we fluorescently-labelled Ras-related nuclear protein Ran, nucleoporin Nup62 or histone H2B (Fig. 1a) as nuclear size markers for live cell fluorescent microscopy. Cell size decreased during hyperosmotic stress and increased during hypoosmotic stress (Fig. 1a). Along with the cells, nuclei shrink and deform, or swell and round up (Fig. 1a). To determine the timescale of these nuclear size changes, we combined live cell fluorescent confocal microscopy and microfluidics, as this approach allowed continuous imaging of cells under controlled changes in osmotic conditions (Fig. 1 b, c, Supplement Fig. 1, Supplement Movies 1-4). Upon osmotic stress, both cell and nuclear size changed in a second to minute timescale. From the time-lapse confocal stacks, we segmented the cellular and nuclear volume and determined that the average cellular size decreased to 0.7-fold of its initial size, corresponding to a reduction of ∼180 μm^3^ within ∼3 min during hyperosmotic stress (n = 12 cells for control and n = 9 for hyper OS) (Fig. 1b, Supplement Fig. 1a). The nuclear volume similarly decreased on average to 0.7-fold of its initial size, a change of approximately 4.3 μm^3^ within ∼50 sec of hyperosmotic stress (n = 9 cells for both control and hyper OS) (Fig. 1c, Supplement Fig. 1b). Although both cytosolic and nuclear volumes decreased by a similar ratio, the whole cell responded slightly earlier to osmotic stress. The adaptation of the nuclear volume initially lagged behind but subsequently caught up at a slightly faster rate. During hypoosmotic stress, the average cell size increased 1.7-fold, corresponding to a ∼410 μm^3^ increase within ∼3 min (n = 7 cells for control and n = 6 for hypo OS) (Fig. 1b, Supplement Fig. 1a) and nuclei expanded 1.4-fold of the initial size, corresponding to a ∼6.0 μm^3^ increase within ∼130 seconds (n = 15 cells for control and 17 cells for hypo OS) (Fig. 1c, Supplement Fig. 1b).

**Figure 1:**
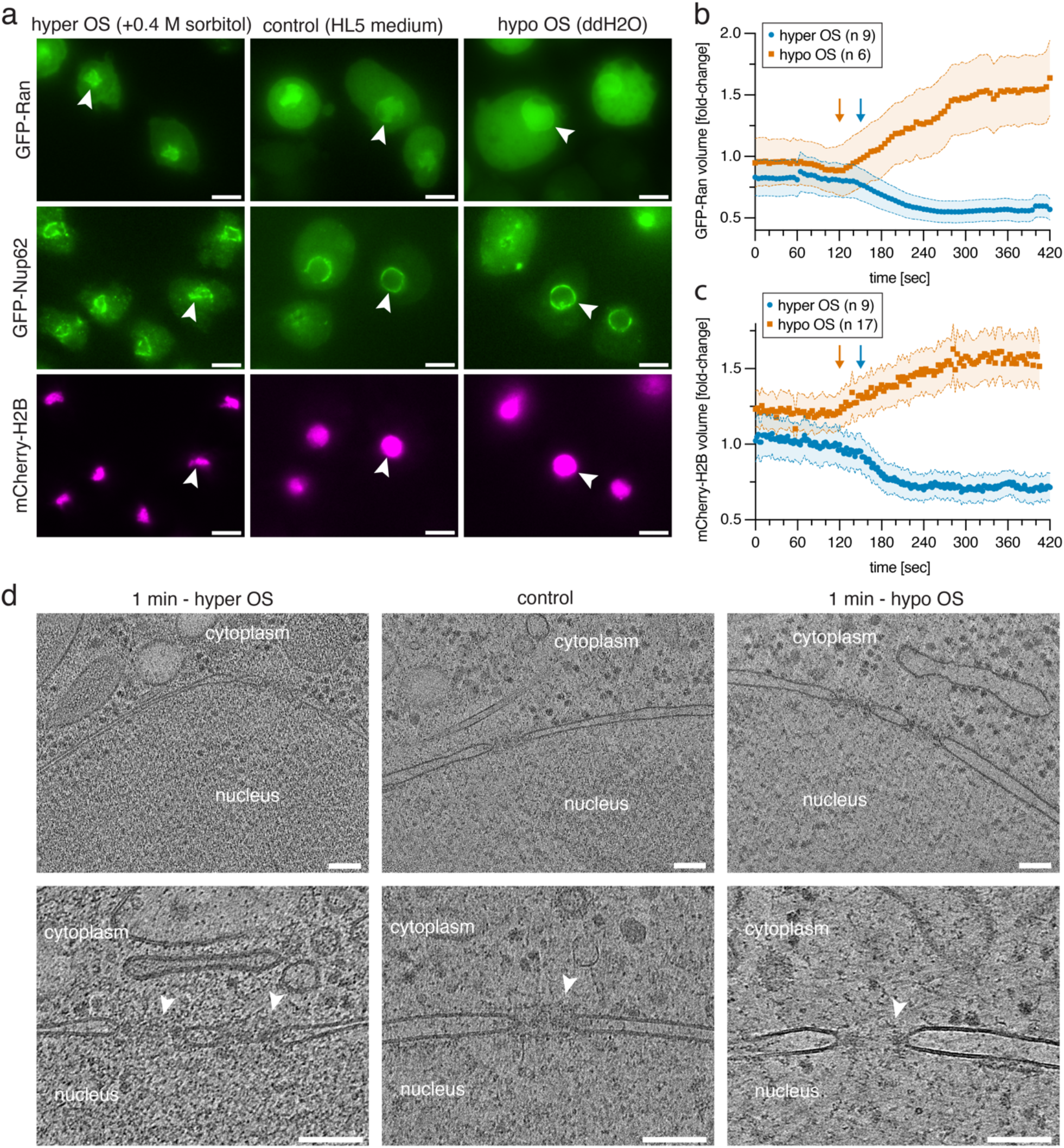
*D. discoideum* cells rapidly adapt to osmotic stress (OS) by changing cellular and nuclear size. **a**, Live cell fluorescence microscopy images of cells overexpressing nuclear marker proteins GFP-Ran (top row), or GFP-Nup62 (center row) and mCherry-H2B (bottom row) in control conditions (center panels) and after ∼3 min of hyperosmotic stress (left panels) or hypoosmotic stress (right panels). Arrowheads highlight one nucleus in each panel. **b**, Fold-change of the cellular Ran-GFP volume from confocal z-stacks during OS normalized to average of control cells as estimate of *D. discoideum* cell volume changes. **c**, Fold-change of mCherry-H2B volume from confocal z-stacks during OS normalized to average of control cells as estimate of *D. discoideum* nuclear size. Standard deviation is shown as shaded error margins. Approximate timepoints of solution exchange (∼150 sec for hyper OS and ∼120 sec for hypo OS) in microfluidics imaging chamber are indicated by blue (hyper) and orange (hypo) arrows in (b) and (c). **d**, Slices of cryo-tomograms at lower and higher magnification of control condition (center panel), and after applying ∼1 min hyper OS (left panel) and ∼1 min hypo OS (right panel). Cytoplasm and nucleus are labeled and white arrowheads indicate nuclear pore complexes in the nuclear envelope. Scale bars: 5 μm in (a), 100 nm in (d).

### Stress-induced cellular volume changes prompt crowding and molecular rearrangements in cyto- and nucleoplasm

Its rapid volume adjustment makes *D. discoideum* an ideal model organism to investigate how molecular crowding or dilution impact on the cell’s macromolecular organization, independently from longer-term transcriptional and proteomic regulation. We thus set out to visualize the cellular ultrastructure and macromolecules at high resolution directly in cells by cryo-ET. For this, we vitrified cells by plunge-freezing approximately one minute after the application of hyper- or hypoosmotic stress (Supplement Fig. 2a). We then prepared cryo-focused ion beam (cryo-FIB) lamellae and targeted areas in the nuclear periphery for cryo-ET in order to capture both nuclear and cytoplasmic regions (Fig. 1d, Supplement Fig. 2b,c). We compared these conditions to a control dataset in which cells were grown in HL5 medium (referred to as control medium), prior to plunge freezing ^53^. The tomographic data acquired after ∼1 min of osmotic stress revealed clear differences in the cellular ultrastructure. While the cellular components and macromolecules in cyto- and nucleoplasm were more densely packed during hyperosmotic stress, the contents appeared less crowded after hypoosmotic stress (Fig. 1d, Supplement Movies 5-7). To quantify the effects of osmotically induced volume change on cytoplasmic protein concentrations and to quantify molecular crowding overall, we used ribosomes as a point of reference, as they are highly abundant and clearly identifiable in cryo-ET tomograms. Using a template matching approach ^54^ with a previously published *D. discoideum* ribosome structure ^53^ (EMD-15808) as the template (Fig. 2), we verified the positions of both cytosolic and membrane-bound ribosomes by classification and eliminated false positive ribosome positions ^53^. We used ribosome numbers as a quantitative measure for cytosolic concentration changes (Fig. 2b) and defined the number of ribosomes within a 100 nm box size (volume corresponds to 0.001 μm^3^) around each ribosome position as an estimate for local cytosolic concentration (Fig. 2c). The mean number of ribosomes per sub volume was 9.2 (51990 sub volumes) under hyperosmotic stress, 6.2 (33589 sub volumes) in control cells and 6.0 (25830 sub volumes) under hypoosmotic stress. As a result, the mean local concentration of ribosomes increased by about 50 % from 10 μM to 15 μM between control and hyperosmotic stress condition. For hyperosmotic stress, the determined ribosome concentration changes correlate well with the overall changes in cellular volume. Classification of ribosomes into different states of the elongation cycle, as previously described for *D. discoideum* and other cell types ^53,55–57^, revealed that translation in general continued under stress conditions, as the most populated states were present in all conditions of *D. discoideum* cells and the largest proportion of states had P-site tRNA present despite changes in cytoplasmic ribosome concentration (Supplement Fig. 3) ^53^. Some re-equilibration between the different states of the translation elongation cycle was however apparent (Supplement Fig. 3).

**Figure 2:**
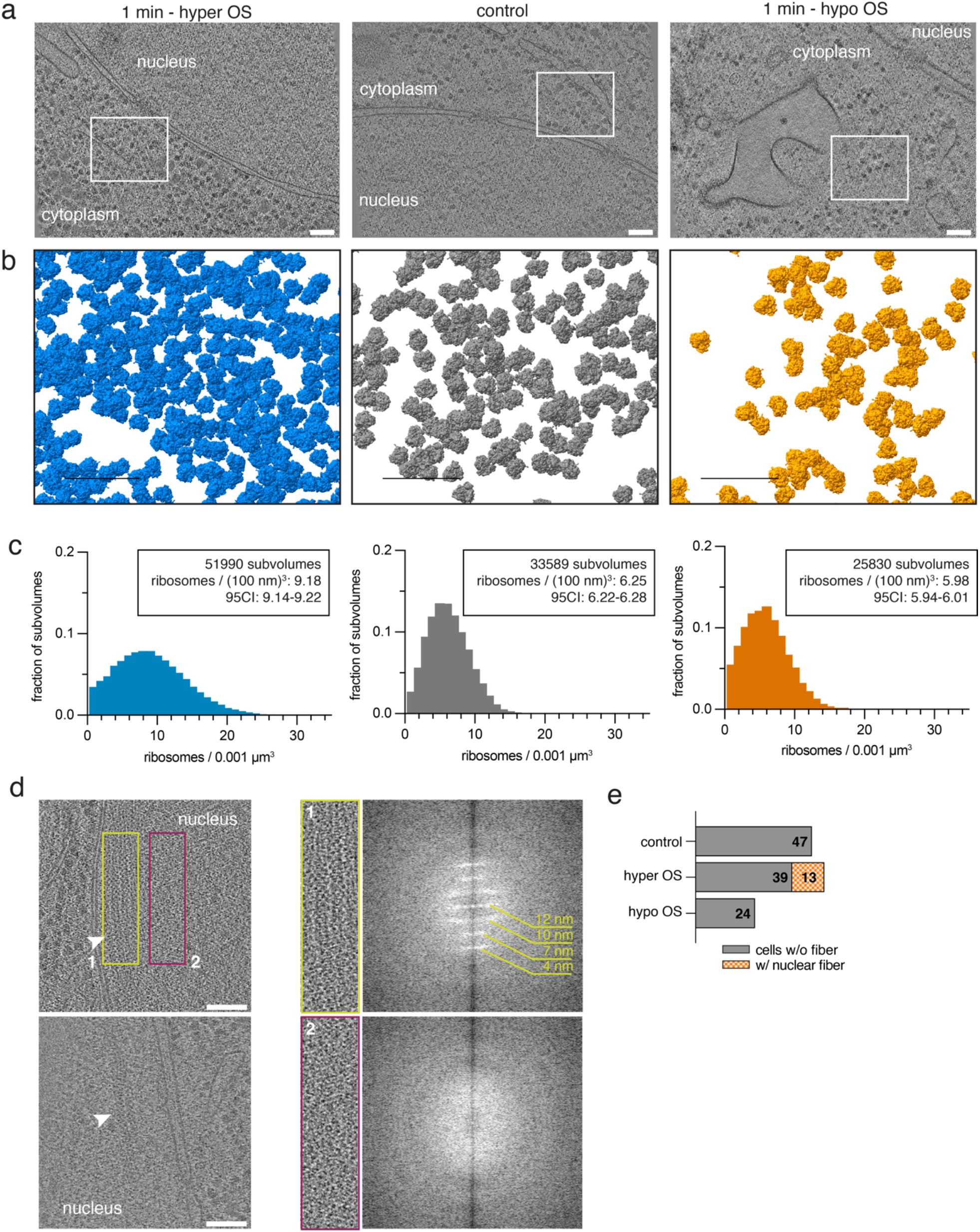
Acute osmotic stress affects molecular crowding in both cytoplasm and nucleoplasm. **a**, Cryo-tomogram slice of cell after ∼1 min of hyper OS, in control conditions and after ∼1 min of hypo OS. **b**, Overview of ribosome positions determined by template matching in a cytoplasmic area (area indicated as white box in panel a). **c**, Histogram of the mean number of ribosomes in cytosolic sub volumes of (100 nm)^3^ around each identified ribosome position. The mean number and the 95 percent confidence interval (CI) are noted for each condition. **d**, Hyper OS led to fiber arrangement of nuclear contents in a subset of cells. Left panels show two tomographic slices with nuclear fibers (White arrows point to fibers). Right panels show zoom of a nuclear fiber (1, yellow) and the control area next to the fiber (2, purple) in upper left panel, as well as the corresponding Fourier transforms. **e**, Fraction of cells with fibers in nucleoplasm longer than 100 nm. 13 of 52 cells during hyper OS had visible fibers in at least one tomogram of their nuclei. No long fibers were detected in nuclei of control cells and cells after hypo OS (47 cells and 24 cells respectively). Scale bars: 100 nm in (a), (b) and (c).

The increase in molecular crowding was also evident in the cell nucleus, where a fraction of nuclei (in 13 out of 52 cells) was found to contain fiber-like arrangements of its nuclear content with an extent of more than 100 nm (Fig. 2d,e). While some repetitive features of this arrangement approximately matched the dimension of nucleosomes (10-12 nm x 4-6 nm) ^58^, we were not able to obtain a conclusive subtomogram average of this structure from our dataset.

### The *D. discoideum* NPC architecture has a unique nuclear ring arrangement with a triple Y-complex configuration

To reveal how nuclear pores adapt to changing molecular crowding conditions and to flow across the central channel, we determined the *in situ* structure of the *D. discoideum* NPC using subtomogram averaging (STA) and integrative modeling of conserved architectural features. A total of 743 NPCs from all conditions combined were analyzed using STA based on previous processing workflows (Fig. 3a, Supplement Fig. 4) (see Methods). We identified yet non-annotated homologues using bioinformatical tools (Supplement Table 2) and built structural models for *D. discoideum* Nups and their interfaces using AlphaFold ^59,60^ (Supplement Fig. 5, Supplement Table 3). We used models of the *D. discoideum* Nups to search for similar folds in PDB using Foldseek ^61^ to validate that the predicted *D. discoideum* Nups have the same fold as the already known Nups (Supplement Fig. 6, Supplement Table 4). Fitting of *D. discoideum* Nup subcomplexes into the respective cryo-EM map allowed us to confidently assign the structured Nups to the overall scaffold (Fig. 3b, Fig. 4a). This revealed an unexpected and unique structural arrangement in comparison to previously published NPCs from other species such as human, *S. pombe, S. cerevisiae, X. laevis* or *C. reinhardtii* ^34–38,47,62,63^.

**Figure 3:**
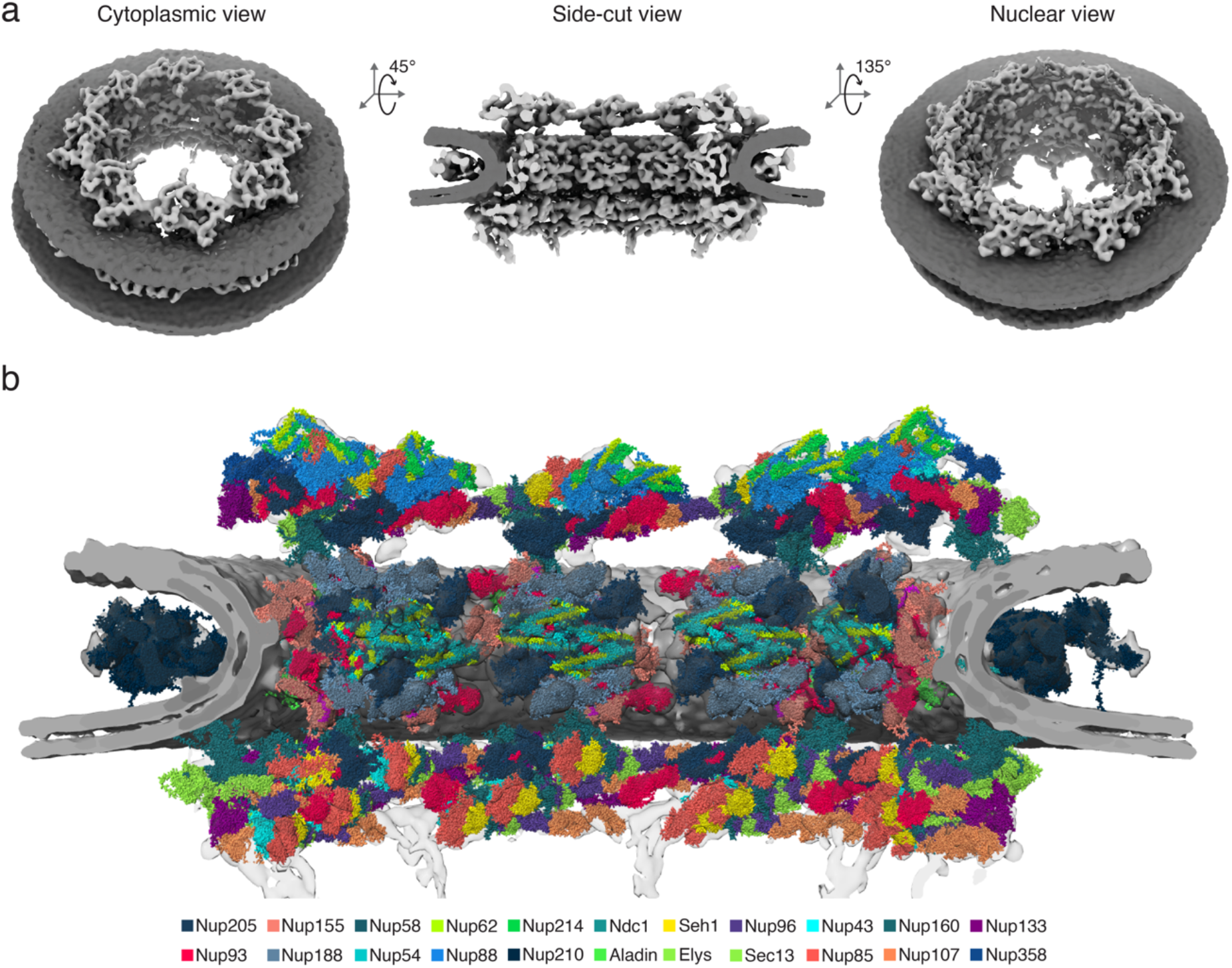
*In situ* structure of the *D. discoideum* NPC using STA and integrative modeling based on AlphaFold structural models. **a**, Subtomogram average of the *D. discoideum* NPC shown as 8-fold rotational average in cytosolic view (left), in side-cut view (center) and nuclear view (right). The NPC is shown in light grey, the nuclear membrane is shown in dark grey. **b, S**tructural model of the *D. discoideum* NPC scaffold built using AlphaFold models and STA density, and guided by the model of the human NPC ^34^.

**Figure 4:**
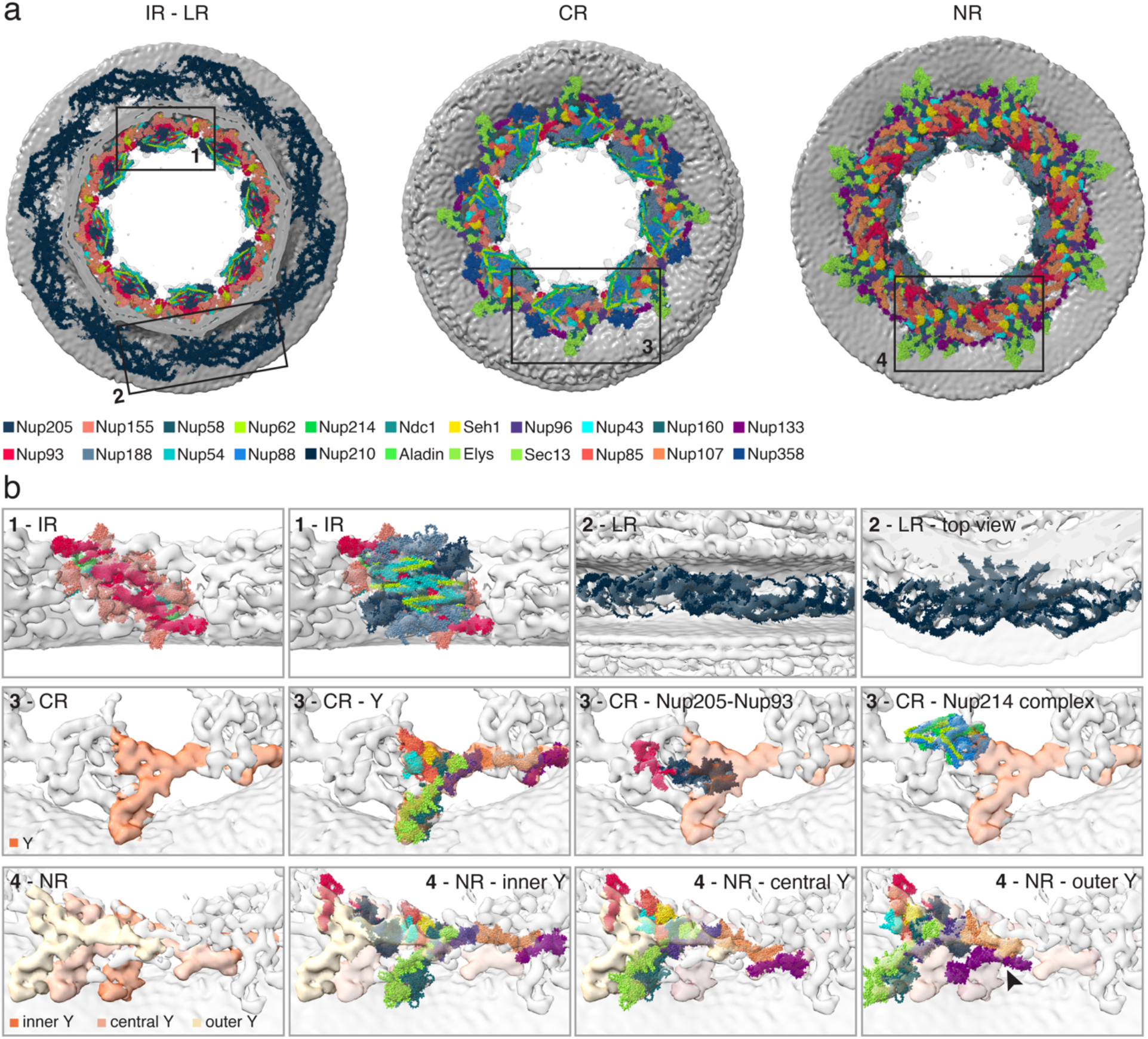
Structural arrangement of the *D. discoideum* NPC. **a**, Structural model overview of (left) the inner ring (IR) and luminal ring (LR), (center) the cytosolic ring (CR) and (right) the nuclear ring (NR). IR and LR are shown as cut view from cytosolic side. **b**, Top left panels: IR spoke shown with predicted AlphaFold models, and only membrane-proximal Nups of IR are shown in the left panel. Top right panels: LR is shown fitted with 8 copies of *D. discoideum* Nup210 homolog per asymmetric unit as side and top view. Center row: CR contains one Y-complex (orange) per structural entity resulting in a concentric ring with 8 copies. Highlighted are the Y-complex, Nup205-Nup93 and two copies of the Nup214 complex with their predicted AlphaFold models. Bottom row: Three Y-complexes form the structural building block of each NR. One copy is located close to the nuclear membrane (orange), one in the center (light orange) and a shorter outer Y-complex (yellow) with non-canonical conformation of Nup133 (indicated by arrowhead; see Supplement Fig. 7 b,c). Each Y-complex and one copy of Nup205-Nup93 are shown fitted with AlphaFold models.

The inner ring (IR) appeared to be conserved in comparison to other organisms (Fig. 4b, panels for 1). Similarly, we observed density for a continuous luminal ring (LR), which fitted to the structural model of eight copies of the *D. discoideum* Nup210 homolog per spoke (Fig. 4b, panels for 2). As for *C. reinhardtii* and *S. cerevisiae*, but in contrast to the human NPC, the cytosolic ring (CR) was composed of a single ring of eight Y-complexes (Fig. 4b, panels for 3). As expected, this ring is further decorated with density reminiscent of one copy of the Nup205-Nup93 complex, two copies of the Nup214 complex and in addition two copies of Nup358 (Supplement Fig. 7a, 8d). In contrast to all organisms for which the NPC was structurally analyzed so far, the nuclear ring (NR) contained three concentric rings of stacked Y-complexes, accounting for a total of 24 copies (Fig. 4b, panels for 4). The inner and the central Y-complex were present in canonical head-to-tail arrangement ^39^. However, the outermost copy of the NR Y-complex was truncated and lacked density at the position where Nup133 would be expected (Fig. 4b, bottom right panel 4, Supplement Fig. 7b). Instead, Nup133 and the interacting C-terminal domain of Nup107 (aa. 708-985) of the outermost NR Y-complex could be fitted in a neighboring yet unassigned density with high confidence (Supplement Fig. 7b, 8a,b,c). This required a conformational rearrangement between the N- and C-terminal domains of Nup107. The presence of the unstructured insertion (aa. 642-702) in the *D. discoideum* Nup107 structured fold enables such conformational rearrangement of the Nup107 (Supplement Fig. 7c). The NR contained one copy of the Nup205-Nup93 heterodimer and nuclear basket attachment for each asymmetric unit (Fig. 4b). To the best of our knowledge, similar architectures of triple Y-complexes have not been reported to date.

### *D. discoideum* NPC scaffolds are resilient to stress-induced diameter changes

Previous work in *S. pombe* has revealed that nuclear pores constrict under conditions of hyperosmotic stress, when nuclear envelope ruffling is observed. This led to the proposal that NE membrane tension regulates NPC diameter ^47^. This model would predict that the relationship between osmotic pressure and NPC diameter should be relevant for both hyper- and hypoosmotic conditions. The latter could not be tested in *S. pombe* cells because their cell wall prevents volume increase. To experimentally test this prediction in *D. discoideum*, we structurally analyzed NPCs after application of hyper- or hypoosmotic stress and compared them to control cells (Fig. 5a,b). The diameter of the *D. discoideum* NPC was on average 72.8 nm (n = 12 grids) in control cells. Indeed, we observed NPC dilation under hypoosmotic conditions, contrasting hyperosmotic stress where NPCs were constricted to 67.8 nm on average, thus confirming our prediction.

**Figure 5:**
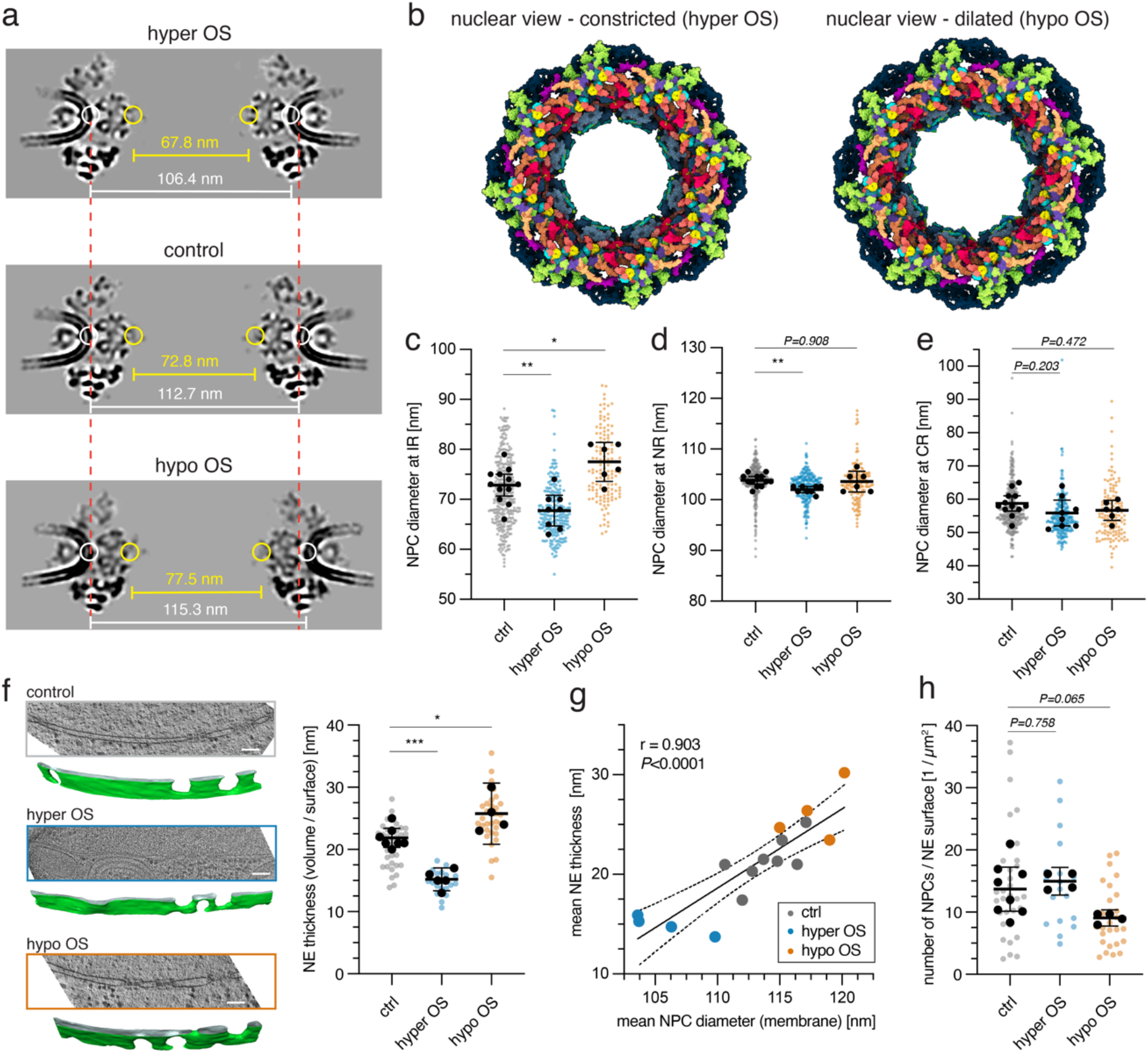
Osmotically-induced constriction and dilation of *D. discoideum* NPCs is coupled to NE thickness, and mediated by movement of nuclear membrane and IR spoke while NR and CR remain rigid. **a**, Comparison of STA maps of constricted NPCs after hyper OS, in control conditions and dilated NPC after hypo OS. The mean diameter between inner ring spoke is shown in yellow (see c for individual measurements), and between outer leaflet of the nuclear membrane in white. The red dashed line indicates the distance of the ONM in the NPC map under control conditions for comparison. **b**, Constricted and dilated NPC model under hyper OS (left) and hypo OS (right). Largest diameter changes are seen for the IR. Nuclear membrane is not shown in the model (Subunit coloring as in Fig. 3b) **c**, NPC diameter measured between IR spokes for individual freezing experiments from STA maps (control: mean diameter = 72.8 nm, 95% CI [70.7, 75.0], n = 12 grids, hyper OS: mean diameter = 67.8 nm, 95% CI [64.7, 70.8], n = 8 grids and hypo OS: mean diameter = 77.5 nm, 95% CI [73.6, 81.4], n = 6 grids). **d**, NPC diameter measured between NR subunits for individual freezing experiments from STA maps (control: mean diameter = 104.3 nm, 95% CI [103.5, 105.0], n = 12 grids, hyper OS: mean diameter = 102.4 nm, 95% CI [101.8, 103.0], n = 8 grids and hypo OS: mean diameter = 104.0 nm, 95% CI [101.9, 106.1], n = 6 grids). **e**, NPC diameter measured between CR subunits for individual freezing experiments from STA maps (control: mean diameter = 58.8 nm, 95% CI [56.5, 52.0], n = 12 grids, hyper OS: mean diameter = 55.9 nm, 95% CI [52.0, 59.8], n = 8 grids and hypo OS: mean diameter = 56.7 nm, 95% CI [53.7, 59.7], n = 6 grids). **f**, Segmentation of NE from tomograms (left) and quantification of NE volume to surface area (right) as approximation for NE thickness in control cells, during hyper OS and hypo OS (control: mean thickness = 21.9 nm, 95% CI [20.3, 23.4], n = 7 grids, hyper OS: mean thickness = 15.2 nm, 95% CI [13.4, 17.0], n = 5 grids and hypo OS: mean thickness = 25.8 nm, 95% CI [20.8, 30.7], n = 4 grids). **g**, NE thickness plotted against the mean NPC diameter measured at the NE membrane for each grid of control cells, during hyper OS and hypo OS shows positive correlation. Spearman correlation was computed with r = 0.903 and *P*<0.0001. The black line represents a linear regression plotted with 95% CI. Data points for controls are shown in grey, hyper OS in blue and hypo OS in orange. **h**, Mean number of NPCs per NE surface area for each grid in control cells, hyper OS and hypo OS cells (control: mean number per μm^2^ = 13.7, 95% CI [10.1, 17.2], n = 8 grids, hyper OS: mean number = 15.0 nm, 95% CI [12.8, 17.2], n = 4 grids and hypo OS: mean number = 9.1 nm, 95% CI [7.8, 10.4], n = 4 grids). Statistical significance was tested using ordinary one-way ANOVA, significance level: * *P*<0.05, ** *P*< 0.01, *** *P*< 0.001, *P* values are stated for non-significant results.

Overall, the *D. discoideum* NPC has a larger diameter as compared to the human NPC (54 nm ^34^) or the *S. pombe* NPC (68.8 nm ^47^), but constriction changes in diameter are notably less pronounced, despite the extreme osmotic changes. NPC constriction and dilation movements were largely performed by the IR spokes, luminal ring and NE membrane (Fig. 5c-e, Supplementary movie 8). While the position of the CR stayed largely fixed (Fig. 5e), the NR underwent a tilting motion and moved closer to the IR spoke during hyperosmotic shock (Fig. 5a, Supplementary movie 9), whereby the NR diameter remained roughly constant (Fig. 5d).

Mathematical modeling had predicted that osmotic pressure across the nuclear membranes and membrane tension should act equivalently. As a consequence, the distance of the inner nuclear membrane (INM) and outer nuclear membrane (ONM) observed by cryo-ET should correlate with NPC diameter, a relationship that was experimentally tested in *S. pombe* cells ^47^. To test if NE thickness and NPC diameter are coupled in *D. discoideum* under conditions of both hyper- and hypoosmotic stress, we systematically quantified the changes to the NE volume in NE segmentations (Fig. 5f). As expected, NE thickness was reduced during hyperosmotic stress, while hypoosmotic stress led to an inflation (Fig. 5f). Indeed, NE thickness and the mean NPC diameter were positively correlated (Spearman correlation coefficient of 0.903) (Fig. 5g). Taken together, these analyses further support a model in which NE tension regulates NPC diameter.

### Pressure-driven volume changes of the nucleus necessitate directional fluid flow across NPCs

The observed fast changes in nuclear volume require directional flow out of and into both, the cell and the nucleus, due to the osmotic gradient. To estimate the flow that occurs across a single nuclear pore, we first estimated the number of NPCs in *D. discoideum* by integrating the STA and NE segmentation data with light microscopic data (see Methods for detail). The average NPC density was 14 NPC / μm^2^ (95% CI between 10-17) in control, 15 NPC / μm^2^ (95% CI between 13-17) in hyperosmotic stress and 9 NPC / μm^2^ (95% CI between 8-10) in hypoosmotic stress conditions (Fig. 5h). From confocal stacks, we determined the mean nuclear volume to ∼20 μm^3^ (N = 50 cells) in control cells, ∼26 μm^3^ (N = 17 cells) during hypoosmotic stress. As a result, *D. discoideum* cells have on average about 380 to 540 NPCs (see Methods). In comparison, this is more than estimated for yeast (100-200 nuclear pores ^64^, but less than in human hepatocytes or in U-2 OS cells (2000 nuclear pores with 14-16 NPCs / μm^2 65^ or 3000 nuclear pores, respectively ^66^).

In an estimate of the actual fluid flow across the nuclear envelope driven by the osmotic imbalance and the resulting changes in cell and nuclear volume, we account both for water passage through the semi-permeable lipid membranes and for flow of solvent including osmolytes through the NPCs. We integrated our experimental data into a mathematical model of hydrodynamics across NPCs to quantify the fluid flow required for nuclear volume adjustments (Fig. 6b). Considerations and simplifying assumptions in our model are laid out in detail in the Methods section. Those are, in brief, an idealized spherical shape for the cell and nucleus, a previously determined water permeability for the biological membranes ^67,68^, and rapid mixing by Fickian diffusion within the cytoplasmic and nuclear compartments. To our assessment, the largest uncertainty is the unknown lipid composition of the nuclear membranes for which we consider the lipid composition of the ER and the experimental studies on their permeabilities, suggesting that it is considerably higher as compared to the plasma membrane (PM) ^68–72^. We assessed the robustness of our model to this uncertainty by considering a range of 35 − 700 *μm* · *s*^−1^ for the NE permeability, referred to as ‘uncertainty range’ below. Taking these considerations into account, four tightly coupled differential equations for the four time-dependent variables (concentrations and radii of cytoplasm and nucleus) were obtained (see Methods section). The differential equations are solved numerically for cells experiencing hyperosmotic (Fig. 6c,d,e) and hypoosmotic (Supplementary Fig. 11) stress.

**Figure 6:**
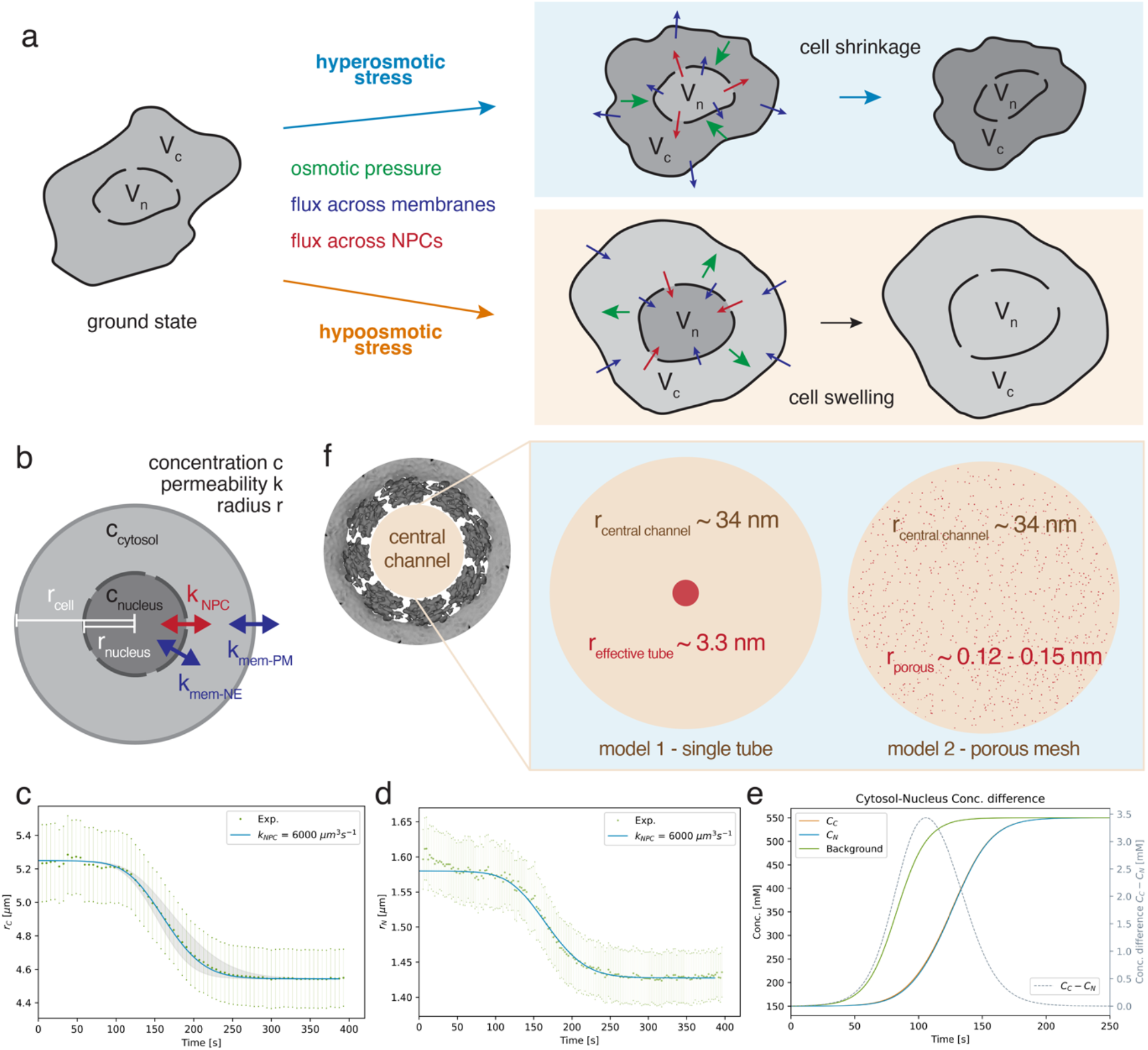
Model for porous flow across nuclear pore complexes. **a**, Schematic of cellular and nuclear volume adjustments during acute osmotic stress. Osmotic pressure causes cell swelling in the case of hypoosmotic stress and cell shrinkage for hyperosmotic stress. Fluid flow to compensate passes over the plasma membrane. Osmotic pressure acts similarly on the nucleus where flow over both the nuclear envelope and through nuclear pores reestablish equilibrium between buffer, cytosol and nucleus. **b**, Schematic model of idealized spherical cell and nucleus assumed for model calculations, with the radii (r) and osmolyte concentrations (c) of cell and nucleus matched to volumetric measurements during osmotic shock. Their time dependence is governed by fluid flow across the PM, the NE and NPCs. **c**,**d**, Experimental and theoretically predicted time dependent cell radius (c) and nucleus radius (d) for hyper OS. Gray lines in C show the theoretical predictions for the PM permeability k_mem-PM_ of 30 *μm* · *s*^−1^ and 40 *μm* · *s*^−1^. **e**, Theoretically predicted time dependent concentrations of cytoplasm and nucleoplasm with experimentally determined buffer concentration for hyper OS. Shown on RHS vertical axis (gray; dashed line) is the difference between the predicted concentrations. **f**, Schematic display of central channel permeability from mathematical models during hyper OS based on experimental data. Left schematic shows the effective channel size radius of ∼3.3 nm if one continuous single path through the FG mesh is assumed according to Hagen-Poiseuille law (model 1). Model 2 shows the FG mesh as homogenous porous medium with a critical radius of ∼0.12-0.15 nm according to Darcy’s law.

Under hyperosmotic stress (Fig. 1b,c, Supplement Fig. 1), we obtained the best fit to the confocal microscopy data for a permeability coefficient of 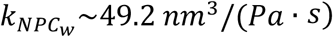 (uncertainty range: 23.1 – 92.3) for one NPC. Translated into fluid flow across a single NPC this equates to ∼4.5 × 10^−4^*μm*^3^*s*^−1^ for a concentration gradient of 0.2 mM across the NE. For reference, the volume contained within a central channel of a single NPC is about 5 × 10^−4^*μm*^3^.

### The permeability barrier remains intact during fluid flow across nuclear pores

The central channel is, however, not empty but densely packed with FG-Nups. Considering the estimated flow rates, we wondered if the permeability barrier for macromolecules also remains intact under acute osmotic stress conditions. Neither under the hyper- nor under hypoosmotic stress conditions, the tomographic data indicated mixing of the observable nuclear and cytosolic features. Neither were ribosomes observed to leak into the nucleus under conditions of hypoosmotic stress nor was chromatin observed to migrate into the cytosol subsequent to hyperosmotic stress. We thus experimentally tested the intactness of the permeability barrier of the nuclear envelope for larger macromolecules, with 500K-FITC-dextran, which did not enter the nucleus following acute nuclear volume increase (Supplement Fig. 10a,b). These observations suggest that the FG-Nup mesh inside the central channel remains intact and, importantly, that *D. discoideum* nuclei did not rupture during nuclear volume changes. Therefore, fluid flow across nuclear pores did not include any larger macromolecular assemblies.

### Porous flow model for fluid flow across nuclear pores during hyperosmotic shock

Having established the NPC permeability coefficient and confirmed that the NE barrier function stays intact, the effective pore size of the NPC can be estimated. Using the Hagen-Poiseuille law for pipe flow, we convert the estimated permeability coefficient of 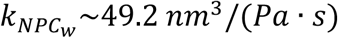 for a single NPC under hyperosmotic conditions into an effective radius of ∼ 3.3 nm (uncertainty range: 2.8 – 3.9 nm) for a single ∼100-nm long tube and a solvent viscosity *μ* about 10-times that of water ^73,74^.

Models of flow through a porous medium based on Darcy’s law ^75–79^ offer a more realistic description of fluid passage through the dynamic network of FG-Nups in the central channel of NPCs ^31,80–82^. We can then relate the permeability coefficient of 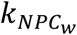 to the porous medium permeability coefficient *k*_*Darcy*_ (Methods section), and in turn estimate an effective radius of the fluid pores using the NPC geometry and the solvent fraction in the pore as additional inputs. For hyperosmotic stress, we obtain *k*_*Darcy*_∼0.012 *nm*^2^ (uncertainty range: 0.006 – 0.024 nm^2^), resulting in a critical radius *r*_*CR*_ of flow tubules ranging from ∼0.12 – 0.15 nm (uncertainty range: 0.08 – 0.22 nm) for porous channels in the homogenous media, wide enough for water passage (Fig. 6f, right panel).

### Flow is larger during hypoosmotic shock as compared to hyperosmotic shock

Under hypoosmotic stress, modelling is complicated by the buildup of tension in the nuclear envelope caused by swelling of the nucleus. By setting the tension factor to σ_*NE*_ = 0.05 Nm^−1^, a value obtained by equating the osmotic pressure to the tension-related hydrostatic pressure ^83^, we estimate a permeability coefficient of 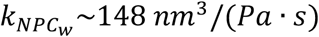 for one NPC during hypoosmotic stress (Supplement Fig. 11). Our results thus indicate ∼3 times larger fluid flow across NPCs compared to hyperosmotic stress. For the Hagen-Poiseuille model with a single cylindrical central opening, we obtain an effective tube radius of ∼4.41 nm (uncertainty range: 2.90−5.55 nm), and for the porous flow model, the effective tubule radius is ∼0.21−0.27 nm (uncertainty range: 0.08-0.40 nm) during hypoosmotic stress (Supplement Fig. 11i). This increased permeability is in accord with the increased NPC diameter (Fig. 5a, b).

## Discussion

Taken together, our analyses point to the following model. In response to osmotic stress, water first flows across the plasma membrane and then across the nuclear envelope. In addition, water and osmolytes pass through nuclear pores. Our mathematical model estimates the effective size of the associated fluid flow, which we then turn into an effective channel opening of the NPC. Over time, the osmolyte concentration is re-equilibrated with adjusted volumes of both cytoplasmic and nuclear compartments. As the permeability barrier of the NPC remains intact during this re-equilibration, our porous flow model appears appropriate to describe the flow within its central channel.

The current view on nucleocytoplasmic transport knows two mechanisms for translocation of molecules across NPCs: passive diffusion and active transport. Passive diffusion allows macromolecules smaller than 30 - 60 kDa to diffuse in equilibrium between the nucleus and cytoplasm while in contrast, active transport is a mechanism that requires binding of a transport receptor to the cargo molecules as well as the RanGTP gradient to facilitate movement through the FG mesh of the central channel against a concentration gradient ^24–26,84^. The concept of porous flow presented here is a functionally distinct translocation mechanism, because it is dependent on physical pressure. Here, cellular solvents directionally flow across the NPC in response to a gradient or force, while the permeability barrier function to large macromolecules remains intact.

We show that volume equilibration processes during osmotic stress in *D. discoideum*, which result in nuclear size adaptations, necessitate considerable fluid flow across the NPC central channel, with flow rates greater than the complete central channel volume each second. At the same time, the central channel is not empty but contains high concentration of FG Nups, cargo and transport factors ^81,85^, additionally restricting free flow of solvent. Our porous flow model predicts the permeability of *D. discoideum* NPCs to cellular solvent, yet the overall largest factor of uncertainty in our model is the permeability of the NE membrane, which needs to be inferred from parameters of other cellular membranes. Aquaporins, ion channels and lipid compositions are unknown contributing factors to NE permeability ^19,86,87^. While the exact NE permeability is not determined, by providing the range of uncertainty, we believe that our model reflects an appropriate estimate for fluid flow and NPC permeability when osmotic stress is applied to *D. discoideum* cells. Our model indicates that the bottleneck for *D. discoideum* cells during the osmotic volume adaptation of cell and nucleus is the flow across the plasma membrane and that nuclear volume adaptation is not limited by flow across the NE. The *D. discoideum* nuclei are thus sufficiently robust to withstand osmotic stress without impairment of the NPC barrier function for large macromolecules. However, in order to efficiently release osmotic stress through fluid flow and to prevent nuclear rupture events, cells likely require to operate within an optimal range of NPC permeability. Deviations from this optimal range could potentially result in altered sensitivity to osmotic stress conditions. Consequently, mutations affecting the number of NPCs, their distribution on the NE, or central channel FG properties ^88–90^, could affect nuclear permeability and alter the cells’ resilience to osmotic stress. Since our proposed mechanism is directly linked to acute nuclear volume adaptations, it is important for cells that are highly adapted to osmotic stress such as *D. discoideum*. In addition, it may play an important role whenever cells undergo severe and rapid nuclear shape changes. Such processes may include migrating cells, infiltrating cancerous cells or cells in developing tissues ^91–94^.

Our structural analysis shows that *D. discoideum* NPCs have an elaborate scaffold structure with three concentric Y-complex rings on the nuclear side and one on the cytosolic side. This scaffold stoichiometry differs from previously assessed organisms ^95^ and adds to the structural diversity of NPCs of various organisms that can assemble from evolutionary conserved Nups into a functional nuclear pore. In contrast to *S. pombe* and human cells ^34,47^, in *D. discoideum* the NPC diameter adaptations are limited to movements of the IR spoke, luminal ring and the nuclear envelope membrane. For *D. discoideum* NPCs, the cytosolic and nuclear ring scaffold structure are remarkably immobile and inflexible to overall constriction and dilation movements. Of notable difference here are the three Y-complex rings of the NR, that may provide additional rigidity to the scaffold thus limiting constriction and dilation. Besides, additional copies of Nup155 that connect the NR and CR to the IR spoke in human and *S. pombe* NPCs could not be detected in the *D. discoideum* scaffold. Alternatively, their presence may be required to translate movements between rings and their absence may confer additional rigidity to the *D. discoideum* NR and CR.

The functional importance of adaptive NPC constriction and dilation is still poorly understood. We corroborate and expand our current model that membrane tension of the NE that is coupled to osmolarity and enforces NPC dilation ^47^. Other studies had shown in mammalian cells, that hypoosmotic stress is associated with a brief spike in membrane tension at least for the plasma membrane and increased cellular volume ^96^. For *D. discoideum*, we are able to induce NPC dilation past its ground state through hypoosmotic stress within the relevant time-window of elevated membrane tension. We conclude that dilation of NPCs by hypoosmotic stress reinforces our previously proposed model that links membrane tension and NPC diameter regulation. The observed dilation contributes to about 20 percent of the change in nuclear surface area. Functionally, it would be conceivable that NPCs can additionally buffer forces on the nuclear envelope, thereby protecting it from rupture and exposure of the nuclear content to the cytosol. This is especially interesting under conditions of mechanical stress for which a recent study showed increased nucleocytoplasmic transport ^52^. Besides affecting nucleocytoplasmic transport, here dilation and constriction may relieve stress on the NE membrane during extreme conditions.

Another intriguing hypothesis is that NPC dilation and constriction may regulate the flow that can occur across the central channel of the NPC. Indeed, using our model of fluid flow across NPCs based on experimental data, we can link lower flow across NPC in hyperosmotic stress to constricted NPCs and higher flow during hypoosmotic stress to dilated NPCs. Thus, the permeability of the central channel to cellular solvents changes with the NPC diameter. Like a self-regulating valve, NPCs may allow for more fluid flow under conditions when nuclei expand under mechanical stress, thus relieving stress. Inversely, constricting NPCs may protect the nuclear volume when tension is alleviated.

However, such a scenario is complicated by multiple parameters and thus its physiological relevance remains to be further tested in the future. The FG-Nup density may not be distributed homogeneously in the central channel. A proposed reduction at its center ^30,31,97,98^ would result in profound effects on central channel permeability. Critical NPC diameters that may allow for central openings in reconstituted synthetic pores range between 60 – 80 nm ^99^ and are compatible with the NPC diameter range we observe *in situ* for *D. discoideum* cells. Inside of cells, however, the changing osmolarity may also affect the local FG-Nup concentration, and possibly their cohesion and molecular condensation behavior, which may severely influence FG arrangement and flow. The NPC permeability in our porous flow model is compatible with the size range of macromolecules that can undergo passive diffusion. ^27^.

Our estimation of the NPC fluid flow is directly based on experimental data, integrating results from live cell fluorescence microscopy into a mathematical model as a complementary approach is to better understand fundamental biophysical processes as they occur in the cells. An effective diameter of the NPC *in situ* that is available for fluid flow is much smaller than the structurally observable diameter underlines the notion that FG-Nups are densely packed inside of cells. We highlight how NPCs provide additional function during nuclear adaptation to stress-generated forces. The proposed porous flow model for fluid flow across NPCs is likely not only important for understanding how cells cope with environmental stress, but broadly applicable to cellular processes involving temporal nuclear shape changes.

## Methods

### Molecular biology and cell culture

The axenic *D. discoideum* strains used in this study were all generated from an Ax2-214 background strain. GFP-Nup62 and GFP-Ran strains carried randomly integrated GFP-Nup62 or GFP-Ran with G418/neomycin resistance cassette ^100,101^. mCherry-H2B was stably integrated at the *act5* locus of Ax2-214 or GFP-Nup62 strains respectively and selected with 50 μg/mL HygromycinB (Sigma Aldrich) using available genetic engineering tools and electroporation protocols for *D. discoideum* cells ^102^. *D. discoideum* cells were grown as described previously ^53^ in HL5 medium (Formedium) with 50 μg/mL ampicillin and additional antibiotic selection with 20 μg/mL geneticin G418 (Sigma Aldrich) or 50 μg/mL HygromycinB (Sigma Aldrich) at 20 ± 2 °C. Cells were kept either as adherent cells in sub-confluent conditions or as suspension culture at cell density between 1 ×10^5^ cells/ml to 4 ×10^6^ cells/ml. Cells were sub-cultured for a maximum of four weeks before re-growing them from cryostocks.

### Fluorescence Microscopy

*D. discoideum* cells were adjusted to a concentration of 2-3 ×10^5^ cells/ml and allowed to adhere to 35 mm dishes with 14 mm cover glass No 1.5 (Mattek) for 2-4 hrs.

Fluorescence images shown in Fig. 1a were acquired using a Nikon Ti2 epifluorescence microscope with a 100x NA=1.49 TIRF objective, equipped with a SOLA light engine (Lumencor), an Orca flash 4.0 LT camera (Hamamatsu), and GFP/FITC and TRITC/CY3 filtercubes (Nikon). The adherent cells on the cover glass were imaged at room temperature. Focal stacks were acquired of cells with a step size of 0.5 μm. Cells were treated for approximately ∼3 min with either HL5 medium + 0.4 M sorbitol or with double distilled water before taking representative images displayed in Fig. 1a. Equatorial slices through the cell’s nuclei are displayed and brightness and contrast of images was adjusted using Fiji ^103^.

Time-lapse imaging of cells was performed at room temperature with a Leica Stellaris 5 microscope equipped with white light laser, with an HC PL APO CS2 63x water objective with NA=1.20 and operated in resonance scanning mode. For initial assessment and testing imaging parameters, cells were adhered to Mattek dishes as described above and osmotically stressed by manually exchanging HL5 medium with either HL5 medium + 0.4 M sorbitol (hyper OS) or with double distilled water (hypo OS). In order to reliably determine nucleus and cell size, *D. discoideum* cells were seeded in CellASIC ONIX M04S-03 microfluidics plates at a density of 1×10^6^ cells/ml. Using the CellASIC ONIX2 microfluidics device cells were either imaged without media exchange (control cells) or with one minute flow of HL5 medium at 34.5 kPa, followed by six minutes flow with respective osmotic treatment solution. In order to monitor buffer exchange, additional buffer exchange experiments for each microfluidics plate were performed again separately with 500K-FITC-dextran (Sigma Aldrich) at a final concentration of 0.2 μM to the treatment solution. All GFP-labeled proteins and fluorescent dextran were excited at 489 nm and emission detected at 500-575 nm, mCherry-labeled H2B was excited at 587 nm and emission detected at 595-670 nm on HyD detectors in bidirectional scanning mode with line averaging. Focal stacks of cells were acquired as 12-bit images with a pixel size of 82.5 nm, z step size of 0.5 μm and z stack sizes of 10-16 μm, resulting in time intervals of 2, 3 or 5 sec between individual stacks. Time-lapse experiments were usually acquired for about 5-10 min. For measuring nuclear permeability of 500K-FITC-dextran during digitonin-induced cell rupture and nuclear volume change, cells adhered to Mattek dishes were exposed to HL5 +0.4 M sorbitol + 20 μg/ml digitonin + 0.2 μM 500K-FITC-dextran while imaging with the settings described above.

### Cell and nuclear size determination

Image segmentation and volume measurements were done in Fiji ^103^. To estimate nuclear size, raw images were cropped to contain one cell of interest, then gaussian filtered with sigma 2 and a threshold with the Default method was applied for segmentation in Fiji ^103^ using the 3D ImageJ Suite ^104^. For cellular size estimation images were additionally filtered with median filter of radius 10 before thresholding with the Otsu method and segmentation in Fiji using 3D manager. Cells or nuclei that moved out of the field of view during the time course of acquisition were excluded from analysis. For monitoring the buffer exchange with 500K-FITC-dextran in the microfluidics chamber, the mean background fluorescence of five squares with 60-pixel box size in empty regions of the raw image was measured over time.

Dextran permeability was directly assessed within the time frame 1 min before cell rupture and 1 min after rupture. As cells randomly ruptured within the field of view after treatment, 10 min time lapse confocal stacks were taken and individual cells upon rupture were analysed. For this, nuclear size was estimated from segmentations of the mCherry-H2B signal as described above with a manual threshold (lower limit of 170) due to the increased background fluorescence due to addition of 500K-FITC-dextran. The nuclear size was determined from segmentations in Fiji ^103^ using the 3D ImageJ Suite ^104^. The mean 500K-FITC-dextran signal of the nucleus and whole cell was measured for a manually selected area covering the equatorial plane of the nucleus or whole cell. All volume estimates and mean grey values were plotted using Prism9 software and to determine shrinkage and expansion, data was fitted using plateau followed by one phase exponential decay function in Prism9.

### Cryo-EM sample preparation

EM support grids (Au grids 200 mesh, carbon or SiO_2_ foil, R2/2 or R1/4, Quantifoil) were glow discharged with a Pelco easiGlow for 90 sec at 15 mA. Exponentially growing cells were adjusted to a concentration of 2 - 3 × 10^5^ cells/ml. A droplet of 100 μl cell suspension was pipetted on the foil side of the grids and cells were allowed to attach to the grids for 2 - 4 hrs. Cells were vitrified by plunge freezing into liquid ethane using a Leica EM GP2 plunger. For treated cells, the media was exchanged for one minute or in few cases for five minutes prior to plunge freezing with either HL5 with 0.4 M sorbitol (hyper OS) or with double distilled water (hypo OS).

Lamellae were prepared by cryo-FIB milling using an Aquilos microscope (Thermo Scientific) as described previously ^53^. In short, samples were coated with an organometallic platinum layer using a gas injection system for ∼10 sec and additionally sputter coated with platinum at 1 kV and 10 mA current for ∼10 sec. SEM imaging was performed with 2 - 10 kV and 13 pA current to monitor the milling progress. Milling was performed stepwise with a gallium ion beam at 30 kV while reducing the current from 500 pA to 30 pA. Final polishing was performed with 30 pA or in some cases with 10 pA current to reach a target lamellae thickness between 130 - 200 nm. Some lamellae were sputter coated with platinum after polishing for 1 or 2 sec at 1 kV and 10 mA. For some of the samples, the rough milling step to a lamella thickness of approximately 600 nm was carried out in an automated workflow using SerialFIB ^105^, while final polishing was then performed manually as described.

### Cryo-ET acquisition

A cryo-ET dataset of *D. discoideum* cell grown in HL5 medium as control condition was acquired in five independent 48-hour microscope sessions from 14 grids from overall 9 independent plunge freezing sessions and with a total of 87 cryo-FIB milled lamellae (Supplementary Table 1). The control dataset is published as ‘dataset 1’ in the context of ribosome translational states *in situ* ^53^ and available through EMPIAR-11845. The hyper OS dataset was acquired in six microscope sessions from 11 grids with a total of 55 cryo-FIB milled lamellae (Supplementary Table 1) and is available through EMPIAR-XXX. The hypo OS dataset was acquired in four microscope sessions from 7 grids with a total of 61 cryo-FIB milled lamellae (Supplementary Table 1) and is available through EMPIAR-XXX. All data were collected at 300 kV on a Titan Krios G2 microscope (Thermo Scientific) equipped with a Gatan BioQuantum-K3 imaging filter in counting mode. For each grid, montaged grid overviews were acquired. Then montages of individual lamellae were taken with 3.9 nm or 2.8 nm pixel size. Tilt series were acquired using SerialEM (version 3.8.1 - version 4.0.1) ^106^ in low dose mode as ∼6K x 4K movies of 10 frames per tilt image, and motion-corrected in SerialEM on-the-fly. The magnification for projection images of 42000x corresponded to a pixel size of 2.176 Å. Tilt series acquisition started from the lamella pretilt of ± 8° and a dose symmetric acquisition scheme ^107^ with 2° increments grouped by 2 was applied, resulting in 59 - 61 projections per tilt series with a constant exposure time and total dose between 132 - 150 e^-^ per Å^2^. The energy slit width was set to 20 eV and the nominal defocus was varied between -2.5 to -5 μm. The initial dose rate on the detector was targeted to be between ∼10 - 20 e^-^/px/sec.

### Tomogram reconstruction

The motion-corrected tilt series were filtered for dose exposure as previously described ^108^ using a Matlab implementation that was adapted for tomographic tilt series ^109^. Poor quality projections were removed after visual inspection. The dose-filtered tilt series were aligned in IMOD (versions 4.10.9 and 4.11.5) ^110^ with patch-tracking using four-fold binning and a patch size of 500-600 pixel with 0.8 patch overlap low frequency roll-off sigma 0.01 and high frequency cutoff radius between 0.07 - 0.2. After initial assessment, contours were broken and tracks that were visually not well aligned with respect to the remaining tracks were removed. Tomograms were reconstructed as back-projected tomograms with SIRT-like filtering of 10 iterations at a binned pixel size of 8.7 Å for initial tomogram inspection. For NPC STA and ribosome template matching, 3D CTF-corrected back-projected tomograms were generated using NovaCTF ^111^. For ribosome averaging and classification, for compatibility with Relion 3.1 ^112^ and M ^113^, the tilt series were reprocessed in Warp ^114^ with the alignment obtained from IMOD.

### Ribosomes template matching

Template matching was done in STOPGAP ^115,116^ using the *D. discoideum* ribosome map (EMD-15808) as template, resampled to a pixel size of 1.306 nm which corresponding to binning factor 6 of the dataset. Template matching was performed on 3D CTF-corrected back-projected tomograms with an angular search of 20°. The top 1000 cross-correlation peaks were extracted from each tomogram. The obtained coordinate and angles for each dataset were converted into Warp-compatible star files using the dynamo2m toolbox ^117^.

### Ribosome classification and ribosome analysis

Classification was performed as previously described ^53^. In short, bin4 (8.704 Å/px) subtomograms were classified in Relion 3.1 ^112^ to filter out non-ribosomal particles such as membranes and other granular structures. This yielded 51990 / 25830 ribosomal particles (hyper OS / hypo OS), which were then again classified to distinguish between cytosolic (50040 particles for hyper OS, 23470 for hypo OS) and membrane-bound 80S ribosomes (1950 particles for hyper OS, 2360 for hypo OS). For all tomograms, we performed manual cleaning of the cytosolic ribosomes in ArtiaX ^118^ resulted in 49667 (hyper OS) and 22999 (hypo OS) cytosolic ribosomes that were used for M refinement and classification. Then, unbinned (2.176 Å/px) subtomograms (of the cytosolic 80S ribosomes were refined in Relion 3.1. The positions were then imported into M ^113^ to perform multi-particle refinement of the tilt series and the ribosome. Geometric and CTF parameters were sequentially refined. This resulted in consensus maps of the cytosolic 80S ribosome after hyper OS (4.5 Å resolution) and hypo OS (5.9 Å resolution). In order to estimate ribosome concentration, for each cytosolic 80S ribosome position, the number of ribosomes contained in a 100 nm large box was determined. For each dataset, the containing ribosome number of each box was displayed in a histogram (Fig. 2 c). Focused classification and state assignment was performed as described ^53^. First, with a smooth shape mask covering the A-, P-, E-site tRNA positions (10 classes, T=4, 35 iterations). The refinements of each class were then subjected to a second round of focused classification (5 classes, T=5, 35 iterations) with a smooth shape mask covering the factor binding site next to the A-site tRNA position. The resulting classes were all refined, and atomic models of 80S ribosomes with elongation factors and tRNAs bound ^119–122^ were rigid body fitted into the refined maps to identify different states. Resulting states and occupancy were compared to the previously published data ^53^.

### NPC particle selection and subtomogram averaging

Positions of NPCs were manually selected as described previously ^63^ and already during visual inspection of tomograms. NPCs that were located close to the lamella border and thus only minimally contained in the lamella volume were not selected for STA.

For NPC STA, 3D CTF-corrected back-projected tomograms were generated using NovaCTF ^111^. Extraction of particles, subtomogram alignment and averaging was in general performed using NovaSTA ^123^ followed by final alignment using STOPGAP ^115^. NPC averaging was performed as described previously ^39,47^. In detail, an initial average of the whole NPC was obtained using NovaSTA. From this, coordinates of the subunits were determined based on 8-fold symmetry. These positions were used to obtain the average structure of the asymmetric unit of the NPC. The extracted asymmetric units were used for further alignment using a spherical mask covering the whole unit. At this point, the particle positions that were generated based on 8-fold symmetry but not contained in the lamella volume, were manually cleaned from the particle list using the ArtiaX plugin ^118^ for ChimeraX ^124^. Initially all datasets were combined to obtain the overall *D. discoideum* NPC structure. Then averaging was again performed for each dataset separately (control, hyper OS and hypo OS) to generate low resolution maps, focused maps and composite NPC maps for each condition. For subtomogram averaging of individual rings, particle positions were re-centered to the respective area (CR, IR, NR, LR, basket connection at NR, basket filaments) based on their position in low resolution maps. STA was further carried out for individual rings with 4- or 2-fold binned subtomograms and elliptical masks around each area of interest in NovaSTA ^123^. Final alignment was carried out using STOPGAP ^115^. The individual ring averages were b-factor sharpened empirically and low-pass filtered. The individual ring maps used were determined at 4-fold binning, as 2-fold binning did not result in better resolved maps based on FSC evaluation and visual inspection. To generate 8-fold symmetric composite NPC maps, the final averages of individual ring maps were fitted to the lower resolution asymmetric subunit average. The density value of all individual ring maps was adjusted to values to represent features of the composite average at a common threshold level. The 8-fold symmetric composite was then produced by applying symmetry based on the coordinates used for splitting the initial NPC average into asymmetric units.

### NPC diameter measurements

NPC diameters were measured based on the coordinates obtained from STA maps of individual rings using previously published MATLAB scripts and as described previously ^47^. Using IMOD, the features of interest in the STA (e.g., the outer nuclear membrane) were identified in map and coordinates were offset by the shift between the center and the feature of interest.

Only NPCs with more than five subunits were considered for diameter measurement of NPCs. As previously described, for each individual NPC, vectors were determined that connected opposing subunits. The point to which the distance of all vectors is minimal was defined as the NPC center, from which the distance to each subunit was reported as the NPC radius and the NPC diameter for each subunit was calculated. Each grid, since frozen and treated separately, was thus treated as one replicate. For each grid, the mean NPC diameter was calculated (black data points), when more than three individual NPC measurements per grid were possible (Fig. 5c-e). Individual NPC measurements were plotted as smaller colored data points. Data was plotted using Prism9 software. Statistical significance was tested using ordinary one-way ANOVA in Prism9.

### NE segmentation, NPC diameter correlation and NPC density calculation

3D CTF-corrected tomograms as described above at binning factor 8 with a pixel size of 1.74 nm were filtered in IMOD using the SIRT-like filtering option. Subsequently, the NE lumen including its membranes was manually segmented from tomograms using the Amira-Avizo software in approximately every 10 slices and then interpolated. Tomograms with limited visibility of the NE membrane, either as result of osmotic treatment or because of the tilted membrane orientation towards the lamella plane, were excluded from segmentation analysis. For segmentation, we used 34 tomograms from 7 grids for control, 21 tomograms from 4 grids for hyper OS and 34 tomograms from 4 grids for hypo OS conditions. For each segmented NE volume, a surface model was generated. The volume comprising the NE lumen, was normalized to the mean NE surface area of the INM and ONM of the segmented NE volume. This ratio served as an approximation of the mean NE thickness in each tomogram. For each grid, the mean NE volume to surface area ratio (black data points), and individual tomogram values (smaller colored data points) were plotted in Prism9 (Fig. 5f). Statistical significance was tested using ordinary one-way ANOVA in Prism9. By integrating the STA NPC and NE segmentation data, we calculated the density of asymmetric NPC subunits per NE surface area, which is displayed as number of NPC per μm^2^ assuming 8-fold symmetry of the NPCs (Fig. 5h). For confocal stacks of control cells and cells during hypoosmotic stress we estimated an average nuclear volume of 20.1 μm^3^ (SD = 6.3, N = 50 cells) and 26.0 μm^3^ (SD = 8.0, N = 17 cells) respectively. Assuming spherical geometry of the nucleus, this resulted in a surface area of 35.7 μm^2^ and 42.4 μm^2^ respectively. Using the average surface of the nuclear envelope, we extrapolated the NPC number per *D. discoideum* nucleus to between 380 – 540.

### Bioinformatics identification of *D. discoideum* Nups

The Nups have been identified by a sequence profile search against the *D. discoideum* proteome, followed by validation with reverse sequence searches and structure prediction. To build the query profile, each human Nup was used to retrieve orthologous proteins from the Eggnog database ^125^. To assemble and build multiple sequence alignment (MSA) from the list of proteins, the retrieved protein sequences were aligned using the MAFFT algorithm ^126^ and adjusted, e.g., trimmed to retain conserved domains or motifs but to remove variable regions. The adjusted sequences were re-aligned, using local or global alignment dependent on the presence of conserved elements, again within the MAFFT environment. The resulting MSA was used to create a Hidden Markov Model (HMM) profile using the hmmbuild and hmmconvert tools provided in the HMMER 3.1b2 package ^127^. The hmmsearch tool was then used to search the *D. discoideum* proteome (UNIPROT GCA_000004695.1), and the first two hits with the best E-value were considered potential Nup candidates.

The candidates were then validated by additional analyses to confirm orthology or to discriminate between alternative hits. The analyses included a reverse search using phmmer against proteome sequence databases of humans and other species presenting conserved Nups.

Further orthology assessment was carried out by an HMM-HMM profile comparison using the HHPRED suite and predicted secondary structure similarity comparison with proteins deposited in the Protein Data Bank ^128^, a search for transmembrane domains, and other conserved motifs/domains typical for Nups. The reverse search was also performed by (BLASTp) searches against SwissProt and nr databases ^129,130^. Finally, the sequences were validated by the similarity of resulting AlphaFold models to orthologous Nups. The final Nup sequence assignments are summarized in Supplement Table 2.

### Structural modeling of Nups and NPC subcomplexes

The structures of individual Nups and NPC subcomplexes were modeled using AlphaFold ^59,60^ available through AlphaPulldown ^131^ and Colabfold ^132^. The max_recycles parameter was set between 12 to 48, depending on the subcomplex, to ensure convergence. The following models were generated using AlphaPulldown: Nup54 (aa. 180-440)-Nup58 (aa. 160-350)-Nup62 (aa. 520-709)-Nup93 (aa. 1-110), Nup160 (aa. 1000-1791)-Nup85-Seh1-Nup43, Nup160 (aa. 1120-1791)-Nup96-Sec13, Nup107-Nup96, Nup160 (aa. 1-1200)-Elys (aa. 1-1100), Nup35 (aa. 377-460) homo-dimer, Nup93 (aa. 170-979)-Nup35 (aa. 1-360), Nup107-Nup133 (aa. 490-1206), Aladin-Ndc1, Nup205_N-Nup205_C-Nup93 (aa. 100-170), Nup188-Nup93 (aa. 100-170), Nup155, Nup155 (aa. 1176-1575)-Nup98 (aa. 800-1000), Nup214 (aa. 601-901)-Nup88-Nup62 (aa. 521-709), Nup160 (aa. 1-1430)-Elys (aa. 590-1170)-Nup96 (707-889), Nup210 (aa. 1-500) homo-dimer, Nup210 (aa. 1151-1610), Nup210 (aa. 355-785), Nup210 (aa. 680-1150), Nup210 (aa.1050-1355) and Nup210 (aa. 1355-1825), Nup155 and Nup358 (aa. 1-780) (Supplement Fig. 5).

The quality of the AlphaFold models was assessed using the scores provided by AlphaFold: the predicted local-distance difference test (pLDDT), which predicts the local accuracy, and the Predicted Aligned Error, which assesses the relative orientation of the proteins and protein domains. In addition, the models of the *D. discoideum* Nups were validated used to searching for similar folds in PDB using Foldseek ^61^ to validate that the predicted *D. discoideum* Nups have the same fold as the already known Nups.

### Fitting of atomic structures to cryo-ET maps

To generate the model of the asymmetric unit of the *D. discoideum* NPC, we used the model of the human NPC (PDB 7R5J) ^34^ and the model of the CR of the *X. leavis* NPC (PDB 7VOP) ^133^ as templates. First, we fitted the IR and LR of the human NPC into the map of the IR and LR of the *D. discoideum* NPC. Then we fitted the fragments of the CR of the human and *X. leavis* NPC into the *D. discoideum* CR map and the fragments of the human NR model into the *D. discoideum* NR map. Finally, we superposed AlphaFold models of the *D. discoideum* NPC subcomplexes to the human/ *X. leavis* model, and optimized the fits of the *D. discoideum* NPC subcomplexes into the map of the *D. discoideum* NPC using ChimeraX ^124^. Systematic fitting of the Nup133 (aa. 490-1206), Nup107 (aa. 708-985)-Nup133 (aa. 490-1206) was performed using the full cryo-ET map of the NR (Supplement Fig. 7b, 8a, b, c) and Nup358 (aa. 1-780) using the difference maps of the CR obtained after subtracting the density of Y-complexes (Fig. 7a, 8d).

### Systematic fitting

We used the previously published procedure for systematic fitting ^39,134,135^ to locate the atomic structures in the cryo-ET maps. Before fitting, all the high-resolution structures were filtered to 15 Å. The resulting simulated model maps were subsequently fitted into individual ring segments of cryo-ET maps by global fitting as implemented in UCSF Chimera ^136^ using scripts in Assembline ^137^. All fitting runs were performed using 100,000 random initial placements, with the requirement of at least 60% of the simulated model map to be covered by the cryo-ET density envelope defined at a low threshold. For each fitted model, this procedure resulted in ∼100 to 3,000 fits with nonredundant conformations upon clustering. The cross-correlation about the mean (cam score, equivalent to Pearson correlation) score from UCSF Chimera ^136^ was used as a fitting metric for each atomic structure, similarly to our previously published works. The statistical significance of every fitted model was evaluated as a P-value derived from the cam scores. The calculation of P-values was performed by first transforming the cross-correlation scores to z-scores (Fisher’s z-transform) and centering, from which subsequently two-sided P-values were computed using standard deviation derived from an empirical null distribution [based on all obtained nonredundant fits and fitted using fdrtool ^138^ R-package]. Finally, the P-values were corrected for multiple testing with Benjamini-Hochberg procedure ^139^.

### Modeling of the *D. discoideum* NPC scaffold

To assemble the model of the entire NPC scaffold we used the integrative modeling software Assembline ^137^, which is based on Integrative Modeling Platform (IMP) version 2.15 ^140^ and Python Modeling Interface (PMI) ^141^.

In addition to using models of subcomplexes as rigid bodies for fitting in the modeling, several inter-subunit interfaces (Nup107-Nup133, Nup107-Nup96, Nup160-Nup85) and inter-domain interfaces of Nup210 were restrained by elastic distance network derived from AlphaFold models, overlapping and bridging the already fitted models. During the refinement, the structures were used as rigid-bodies and simultaneously represented at two resolutions: Cα-only representation and a coarse-grained representation, in which 10-residue fragments were represented as a single bead. The Cα-only representation was used for all restraints except for the EM fit restraint.

The structures of individual rings (CR, NR, and IR together with LR) were optimized using the refinement step of Assembline to optimize the fit to the map, minimize steric clashes, and ensure connectivity of the protein linkers. The scoring function for the refinement comprised the EM fit restraint, clash score (SoftSpherePairScore of IMP), connectivity distance between domains neighboring in sequence, and elastic network restraints derived from the subcomplexes modeled with AlphaFold. The final atomic structures were generated based on the refinement models by back-mapping the coarse-grained representation to the original AlphaFold atomic models. The stereochemistry of the final models was optimized using steepest descent minimization in GROMACS ^142^.

The structures of the NPC in the constricted and dilated states were generated by fitting the model of the asymmetric unit of the *D. discoideum* NPC into the EM map of the constricted and dilated NPC and refining using Assembline.

### Mathematical model of fluid flow across the NPC and NE during osmotic stress

In our model of fluid flow across the NPC, we assumed an idealized spherical cell containing a spherical nucleus with radii r_C_ and r_N_, respectively. We assumed each compartment to be well mixed with osmolyte concentrations C_C_, C_N_ and C_O_, respectively, in the cytosol (C), the nucleus (N), and outside (O) of the cell. We set the water permeability of the plasma membrane at a value typical for cells, 35 *μm* · *s*^−1 67,68^ (Fig. 6c). For the more loosely packed NE ^68–72^, we assumed a permeability in the range of 35 − 700 *μm* · *s*^−1^, with a typical value of 350 *μm* · *s*^−1^. We included tension contributions to the pressure in the NE only under hypoosmotic conditions, assuming the NE to be relaxed under hyperosmotic conditions. Certain volume fractions inside the cytosol and the nucleus were excluded from the effective solvent volumes undergoing changes osmolarity during the experiment ^143,144^. We estimated that there are ∼450 NPCs per cell (Fig. 5h).

The osmotic pressure *Π* across a semi-permeable membrane is proportional to the difference in osmolyte concentration Δ*C*,

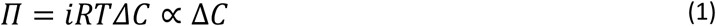

where *i, R*, and *T* denote the van ‘t Hoff index, ideal gas constant and temperature, respectively. The osmotic pressure drives a water flux through the membrane ^145^,

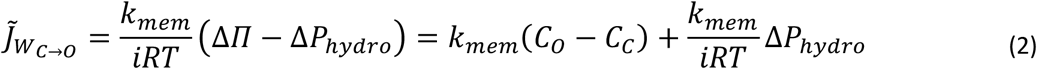

where 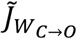 is the number of translocated water molecules (in unit of mole) per unit time and unit area from the cytosol to the outside; *k*_*mem*_ is the membrane water permeability; and Δ*P*_*hydro*_ = *P*_*O*_ – *P*_*C*_ is the difference in the hydrostatic pressure between the nucleus and cytosol. Without hydrostatic pressure change across the plasma membrane, *P*_*O*_ ≈ *P*_*C*_, the flux from the cytosol to the outside (in unit of mole per unit time) is,

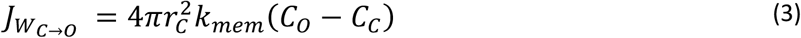

We treat the NE as a single entity, ignoring its small luminal volume. The flux of water from the nucleus to the cytosol then becomes

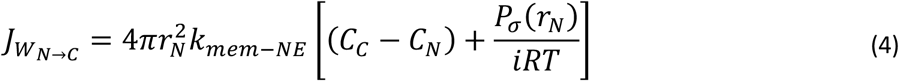

where *k*_*mem*−*NE*_ denotes the overall water permeability across the nuclear envelope and *P*_*σ*_ (*r*_*N*_) (Eq. (5)) denotes the hydrostatic pressure due to the tension in the NE (*σ*_*NE*_), which accumulates as the NE expands. For expanding nuclei, the NE tension cannot be neglected due to the support of its lamina, which provides the NE a larger resistance against expansion compared to the plasma membrane. Note when *C*_*N*_ > *C*_*C*_, the sign of the first term is negative while the second term is positive, hence the NE tension would retard the water diffusing into the nucleus from the cytosol, across the NE. For the pressure created by NE tension we use ^83^

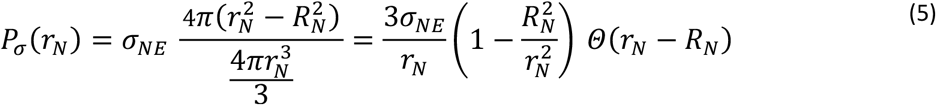

where *R*_*N*_ denotes the radius of the relaxed NE and the Heaviside function *Θ* ensures that tension acts only when the NE is expanded, *r*_*N*_ > *R*_*N*_. The value of *σ*_*NE*_ can be estimated from the experiment by equating the osmotic and tension pressure during the steady state after the hyposmotic shock. Here, *σ*_*NE*_ was estimated to be 0.05Nm^−1^, with a strict upper bound of ≈ 0.1Nm^−1^.

Exchange between the cytosol and the nucleus are further mediated across the NPCs, through which bulk flow (translocation of both the water and solute molecules; 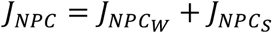), is allowed. Due to the concentration gradient across the NE, *J*_*NPC*_ is driven by both the hydrostatic pressure and chemical potential. Note that the former is partly a consequence of the NE tension, which causes efflux 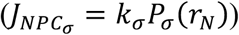 when *r*_*N*_ > *R*_*N*_. Another contribution to the former is the accumulation of the osmotic pressure across the NE (Eq. 1), and the consequent hydrostatic pressure induces the flux from low to high concentration via NPCs, analogous to the “balloon effect”; 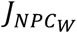 has the same sign as 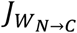 and if we neglect the concentration gradient within the NE, such hydrostatic pressure (due to the build-up of osmotic pressure) is proportional to *C*_*N*_ – *C*_*N*_, ignoring the NE as a separate compartment. In the following, we also ignore diffusive water and solute transport across the NPC, which is overwhelmed by hydrodynamic flow. We also assume that the solute volume fraction is small compared to the water volume fraction, *φ*_*S*_ *≪ φ*_*W*_. Consequently, differential equations for each volume compartment can be formulated based on the expressions for the respective fluxes. In time *dt*, the volume of the cell changes as,

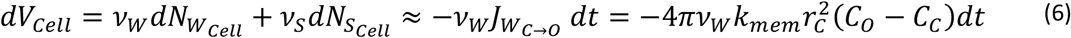

where *ν*_*W*_ and *ν*_*S*_ are the molar volume of water and solute, respectively, and 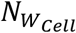 and 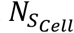 are the total number of water and solute molecules inside the cell (in moles). The volume of the nucleus changes as

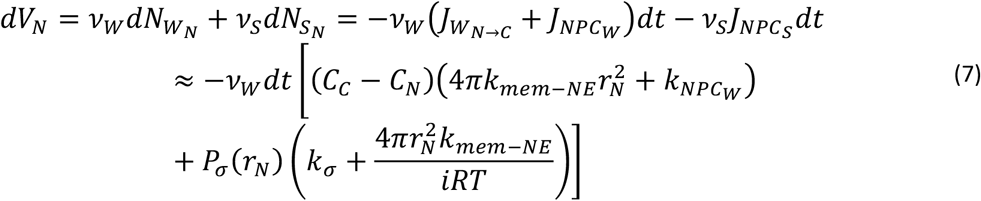

For the solute concentrations in each compartment, we use effective volumes obtained by subtracting excluded volumes (of organelles, genome, nucleoli etc.), which we estimated at 50% of the measured values before the osmotic shock: 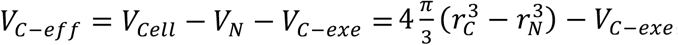, and, 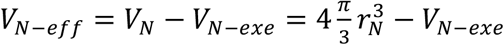. The concentrations at times *t* + *dt* with small *dt* are then

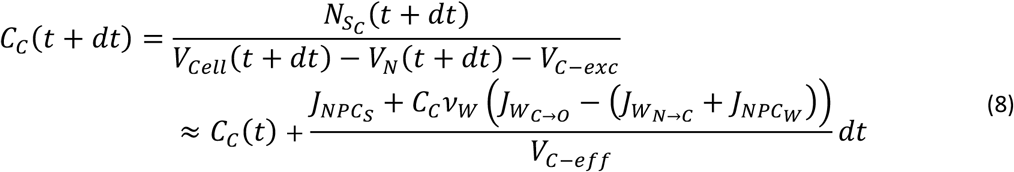

And similarly

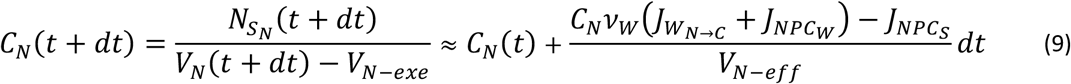

We now rewrite the above expressions as differential equations for the concentrations and the radii of the spherical volumes:

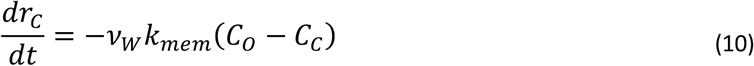

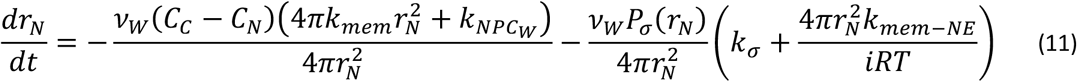

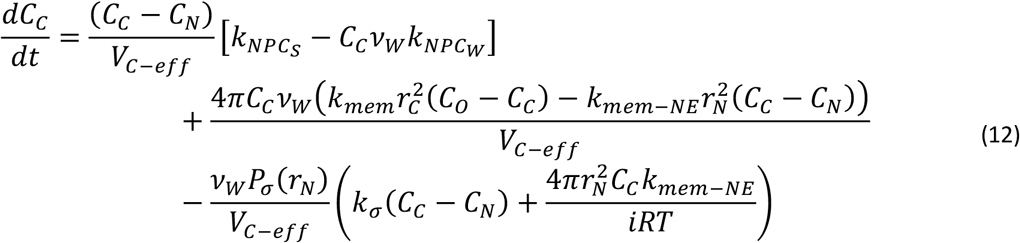

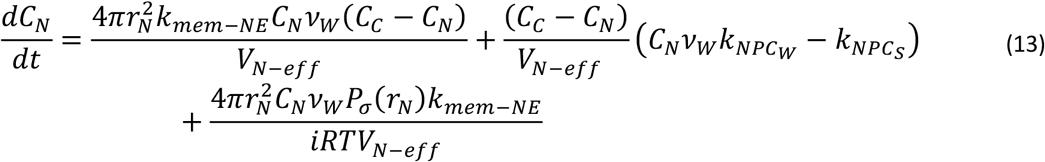

A full list of the constants – together with the values used/estimated in this manuscript – in Eq. (10) – (13) can be found in Supplement Table 4. The differential equations were solved numerically with a Runge Kutta-4 integrator ^146^.

### Porous flow model

Assuming a porous flow model, the fluid flux through the NPC satisfies Darcy’s law ^147,148^,

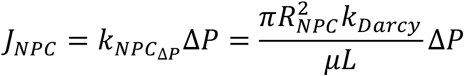

with *μ* the solvent viscosity. For cells undergoing hyperosmotic shock, the pressure difference Δ*P* across the *NE* is due to the accumulation of the osmotic pressure, Eq. (1).

The permeability of the porous medium *k*_*Darcy*_ can be related to critical radius of the tubules through the medium ^75–79^,

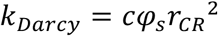

where *c* ≈ 1 is the relative path length and *φ*_*s*_ ≈ 0.5 to 0.9 is the solvent volume fraction in the pore. In our estimates, we set *L* = 100 nm and *μ* to 10 times the water viscosity ^73,74^.

### Uncertainty estimation

Supplement Table. 5 illustrates a complete list of input variables of the model, with the corresponding uncertainty ranges. Based on the second column entry of the input variables, the permeabilities of a single NPC during hypertonic and hypotonic shocks were determined to be 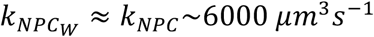 and 18000 *μm*^3^*s*^−1^ (*J*_*NPC*_ = *v*_*W*_*k*_*NPC*_Δ*C*), respectively, by performing a chi square test. The uncertainty in the NPC permeability associated with the uncertainty ranges of the individual non-experimentally predetermined input variables were numerically estimated. Considering that a monotonic change in most of the input variables individually causes a monotonic change in the NPC permeability, the overall uncertainty in the NPC permeability is numerically obtained to be a factor ∼2. However, it is important to note that this seemingly large relative uncertainty is greatly reduced when the permeability *k* is converted into an effective pore radius *r* for Hagen-Poiseuille flow, *r* ∝ *k*^1/4^, or Darcy porous flow, *r* ∝ *k*^1/2^.

Compared to hyperosmotic stress, modelling the hypoosmotic stress is subject to a larger degree of uncertainty and error, especially for later time points. A typical example is illustrated in Supplement Fig. 11a, in which the predicted *r*_*C*_ noticeably deviates from the experimental observation at late times. The major cause of this is the neglection of a plasma membrane tension building up as the cell expands. Relatedly, the chi square test for the hypotonic shock was performed for time up to 300 seconds in Supplement Fig. 11. Furthermore, the modelled NE tension (Eq. (5)) is also subject to a certain degree of uncertainty (*σ*_*NE*_ in Supplement Table 5). The corresponding uncertainty in the NPC permeability was found to be ≈ 10%, which is small compared to the overall uncertainty (∼factor 2).

## Data availability

The NPCs maps reported in this paper will be deposited in the EM Data Bank with accession codes XXX and released upon publication. Ribosome maps will be deposited in the EMDB with accession codes XXX and released upon publication. The composite maps and individual The modeled *D. discoideum* NPC structure will be available at PDB with accession code XXX and released upon publication.

The raw tilt series and alignment files for control conditions, hyperosmotic and hypoosmotic stress conditions will be deposited on EMPIAR with accession codes EMPIAR-11845, XXX and XXX, respectively, and available upon publication. Raw fluorescence data will be deposited and available through the BioStudies database.

## Acknowledgements

We thank Christian Zimmerli, Matteo Allegretti, Max Seidel, Florian Wilfling and Maziar Heidari for advice. We thank Stefanie Böhm for careful reading and advice during manuscript preparation. We thank Sonja Welsch, Mark Linder and the members of the Central Electron Microscopy facility of the Max Planck Institute of Biophysics for technical support and support with data acquisition. We thank Özkan Yildiz, Juan F. Castillo Hernandez and Thomas Hoffmann and the Max Planck Computing and Data Facility for support with scientific computing. H.K. thanks International Max Planck Research School (IMPRS) on Cellular Biophysics and the Max Planck Society.

## Author contributions

P.C.H. conceived the project, designed experiments, performed experiments, analyzed data, and wrote the manuscript. H.K. developed the mathematical model, analyzed data, and wrote the manuscript., A.O.-K. performed structural modeling, analyzed data, and wrote the manuscript., J.P.K. and E.A.-F. performed ribosome classification, analyzed data and edited the manuscript, L.C. performed bioinformatical analysis under supervision of J. K.; both edited the manuscript, B.T. and S.C.L. supported cryo-ET data processing and edited the manuscript, G.H. conceived the project, supervised the project, acquired funding, and wrote the manuscript. M.B. conceived the project, supervised the project, acquired funding, and wrote the manuscript.

