## Supplemental Material for "Nuclear pores as conduits for fluid flow during osmotic stress"

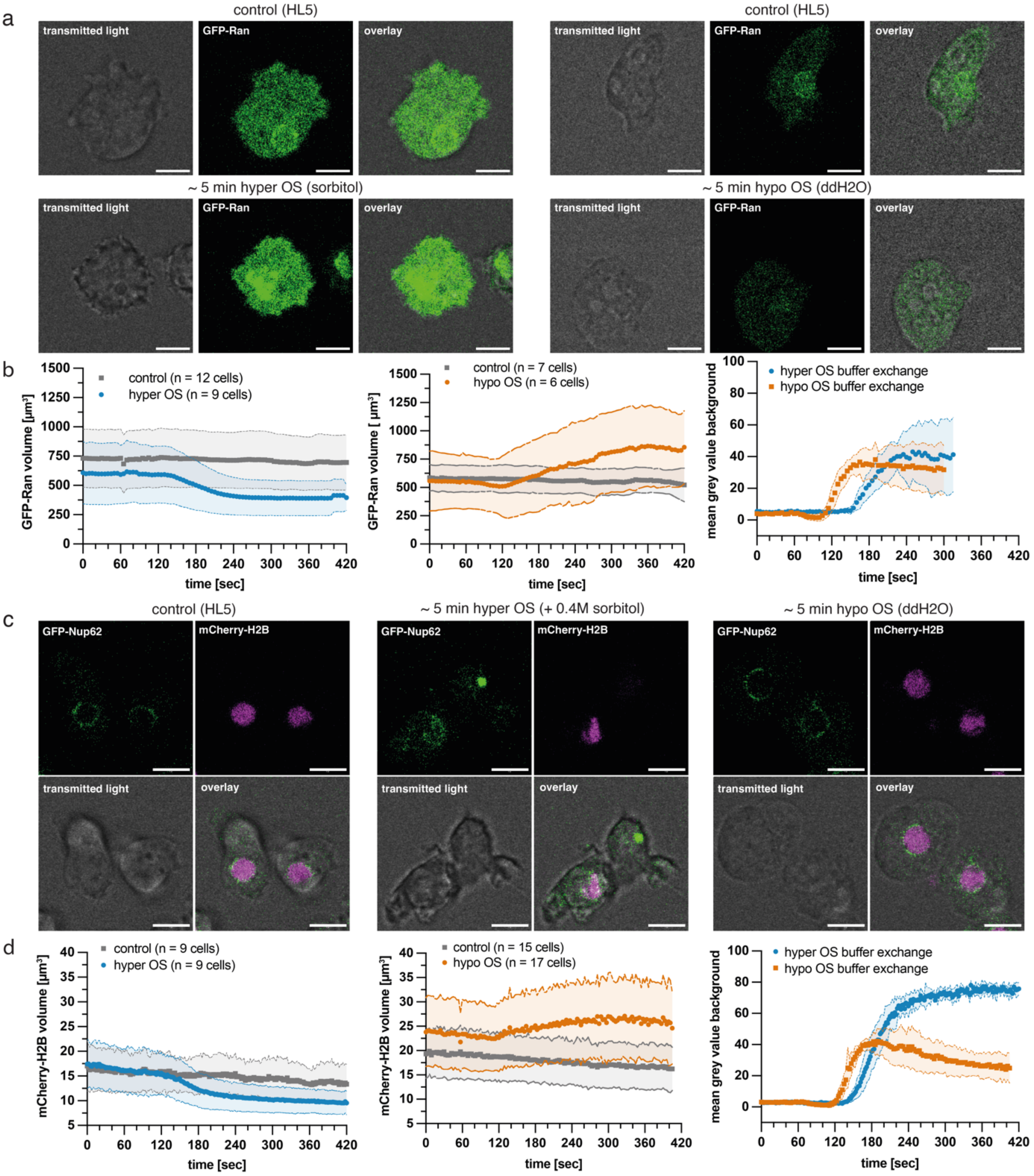


**Supplement Figure 1: Fluorescence microscopy for cell size and nuclear size quantification of osmotically stressed *D. discoideum* cells. a and c,** Live cell images from confocal stacks of *D. discoideum* cells overexpressing GFP-Ran (a) or the nuclear marker proteins GFP-Nup62 and mCherry-H2B (c) under control growth conditions (left panels, grown in HL5 medium), cells after 5 min of hyper OS (center panels, HL5 medium with 0.4 M sorbitol) and cells after 5 min of hypo OS (right panels, $ddH_2O$). GFP Ran and GFP-Nup62 shown in green, mCherry-H2B shown in magenta. **b and d,** Quantification of segmentation of GFP-Ran signal as proxy for cell size (b) or of mCherry signal as proxy for nuclear size (d) in control cells and during hyper OS, and for control cells and during hypo OS. Quantification of background fluorescence to determine the time of buffer exchange with 500K-FITC-dextran in respective osmotic stress condition. Scale bars: 5 μm in (a) and (c). For corresponding time-lapse movie see supplement movies 1-4.

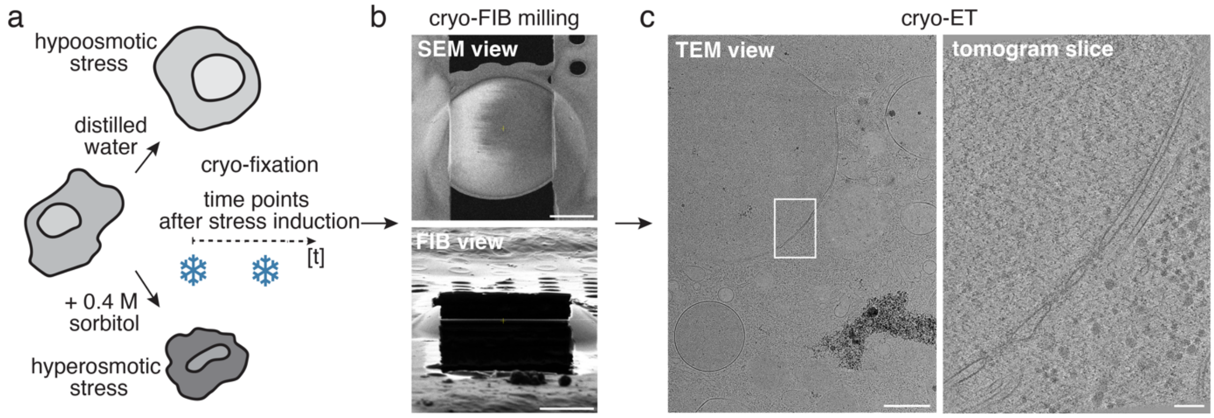


**Supplement Figure 2: Cryo-ET workflow of osmotically stressed *D. discoideum* cells**. **a,** Study design for *in situ* cryo-ET of osmotically stressed *D. discoideum* cells. **b,** Cryo-FIB milling of *D. discoideum* cell, SEM view (top panel) and FIB view of lamella (bottom panel). **c,** TEM overview of lamella area containing nucleus (left panel). Virtual slice through cryo-tomogram (right panel) acquired at the area of the white box. Scale bars: 5 µm in (b), 1 µm in (c) left panel, 100 nm in (c) right panel.

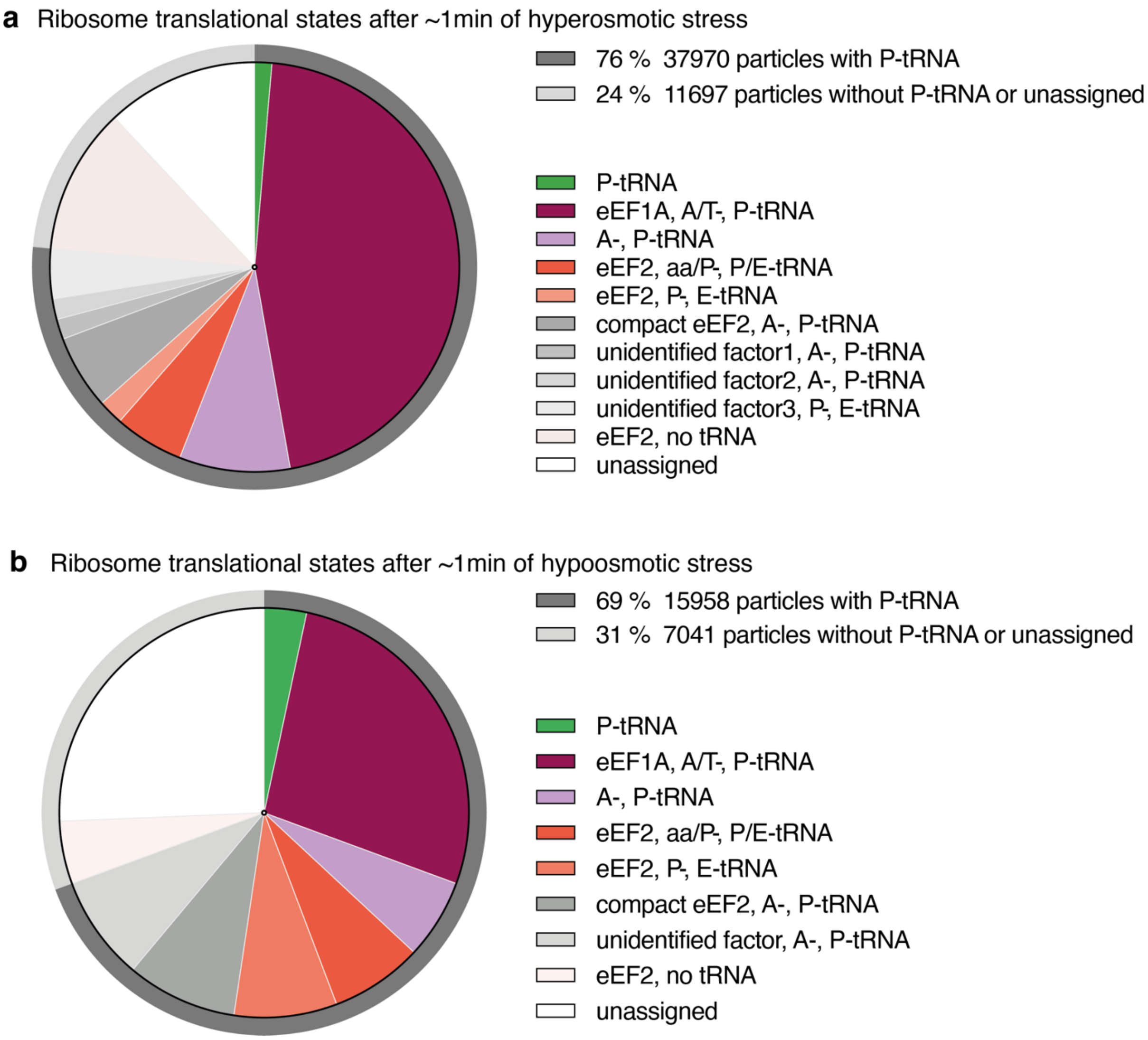


**Supplement Figure 3: Ribosome translational states of osmotically stressed *D. discoideum* cells**. **a,** Translational states of free cytosolic 80S ribosomes in *D. discoideum* cells after 1 min of hyperosmotic stress. 76% of particles have P-site tRNA. **b,** Translational states of free cytosolic ribosomes in *D. discoideum* cells after 1 min of hypoosmotic stress. 69% of particles have P-site tRNA. In comparison, *D. discoideum* grown in HL5 (control condition) had 76% of particles in translating states with P-site tRNAs present and eEF1A, A/T-, P-tRNA as main state [53].

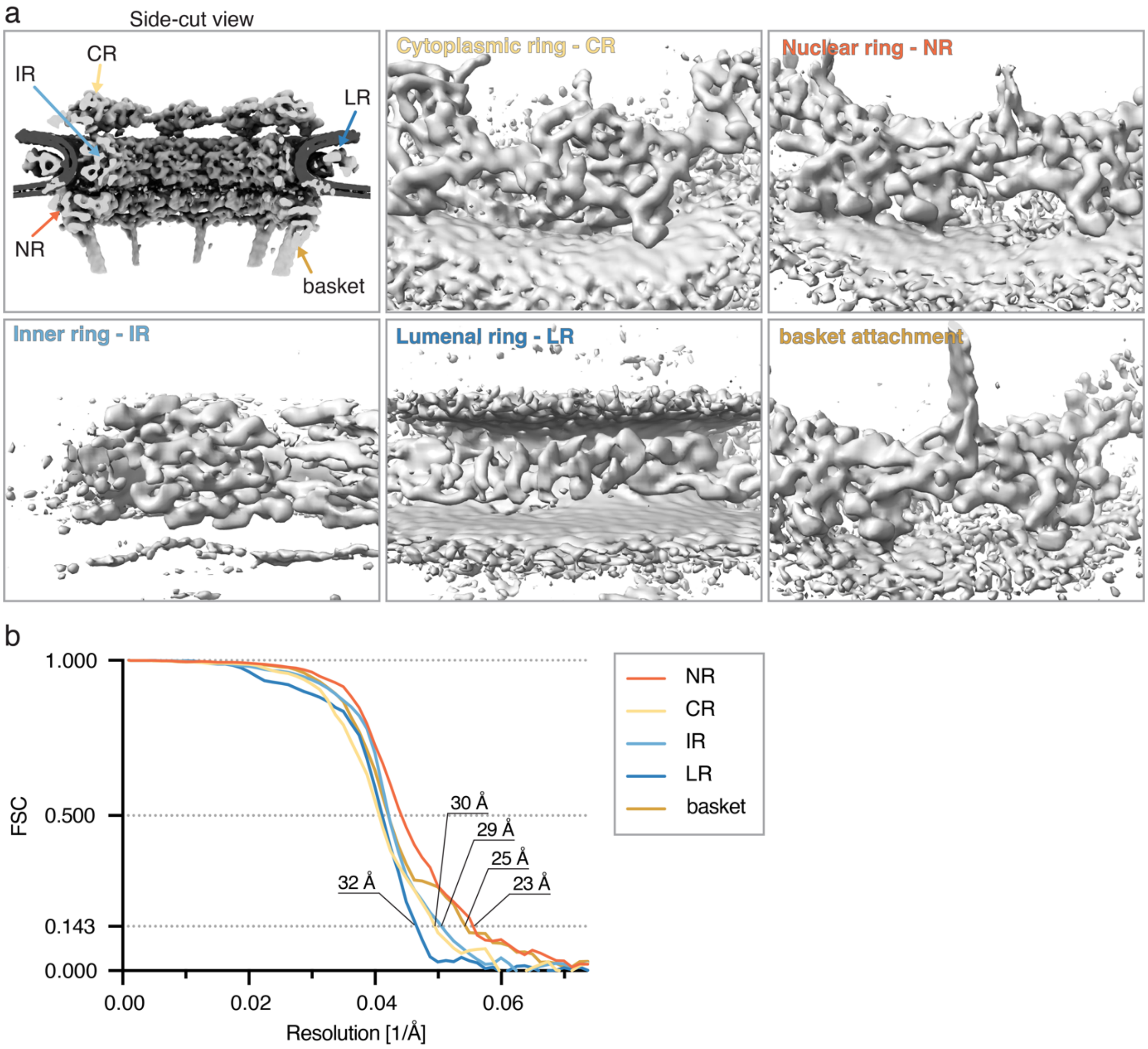


**Supplement Figure 4: STA map of *D. discoideum* nuclear pore complex. a,** Side-cut view of the 8-fold symmetrical nuclear pore structure from a subtomogram average of 4921 subunits, 745 NPCs, from *in situ* tomograms of *D. discoideum* cells grown in control conditions, and during hyper- and hypoosmotic stress combined (top left panel). NPC map is shown in light grey and membrane shown in dark grey. STA maps focused on different regions of the NPC, e.g. CR, IR, NR, LR and region of nuclear basket attachment. **b,** FSC curves of the focused STA maps for individual parts of the NPC, the resolution cut-off at 0.143 is indicated for each focused map shown in (a).

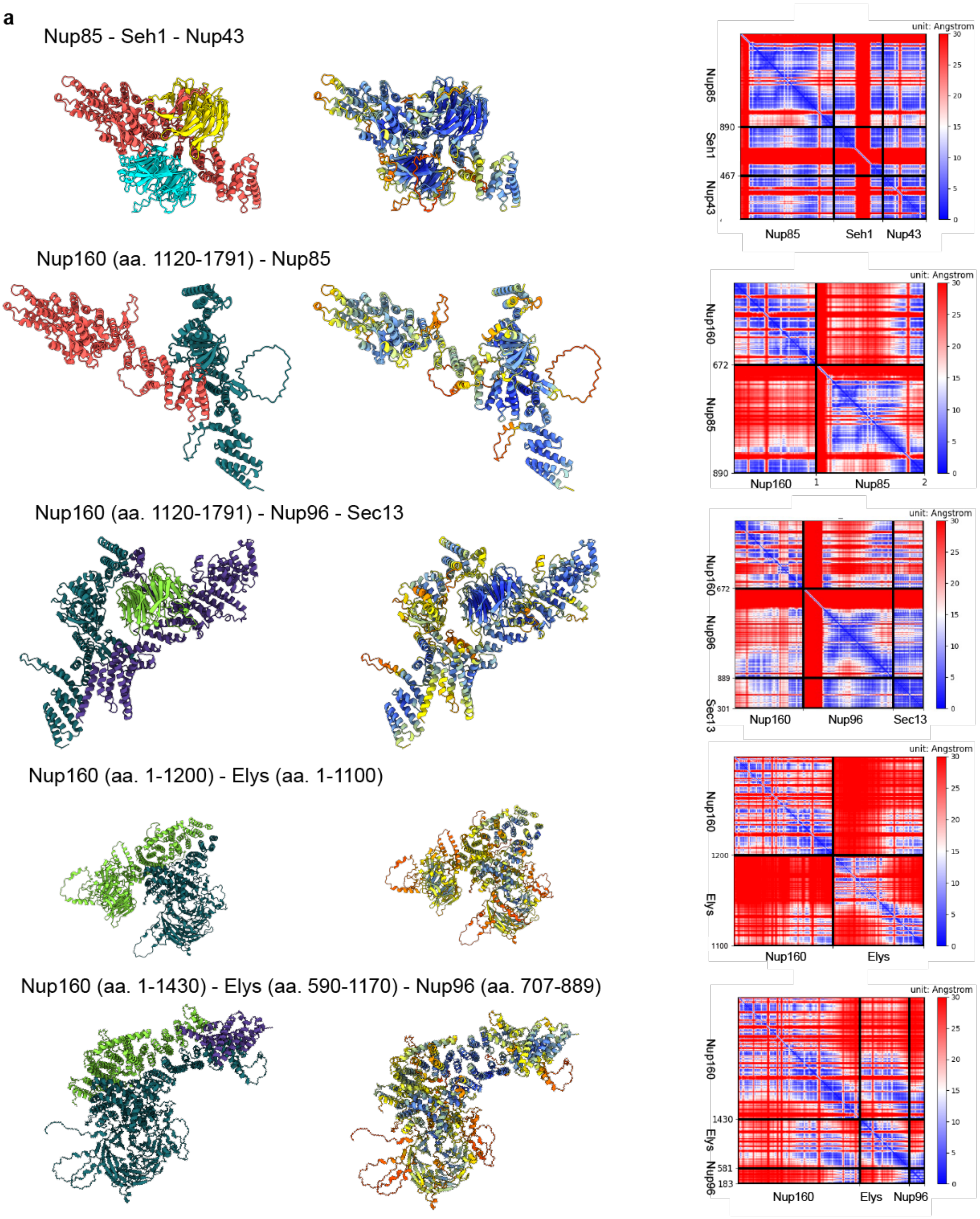
a
Nup85 - Seh1 - Nup43
Nup160 (aa. 1120-1791) - Nup85
Nup160 (aa. 1120-1791) - Nup96 - Sec13
Nup160 (aa. 1-1200) - Elys (aa. 1-1100)
Nup160 (aa. 1-1430) - Elys (aa. 590-1170) - Nup96 (aa. 707-889)
unit: Angstrom
Nup85
Seh1
Nup43
890
467
Nup160
672
Nup96
Sec13
889
301
1200
1100
Elys
1430
581
183

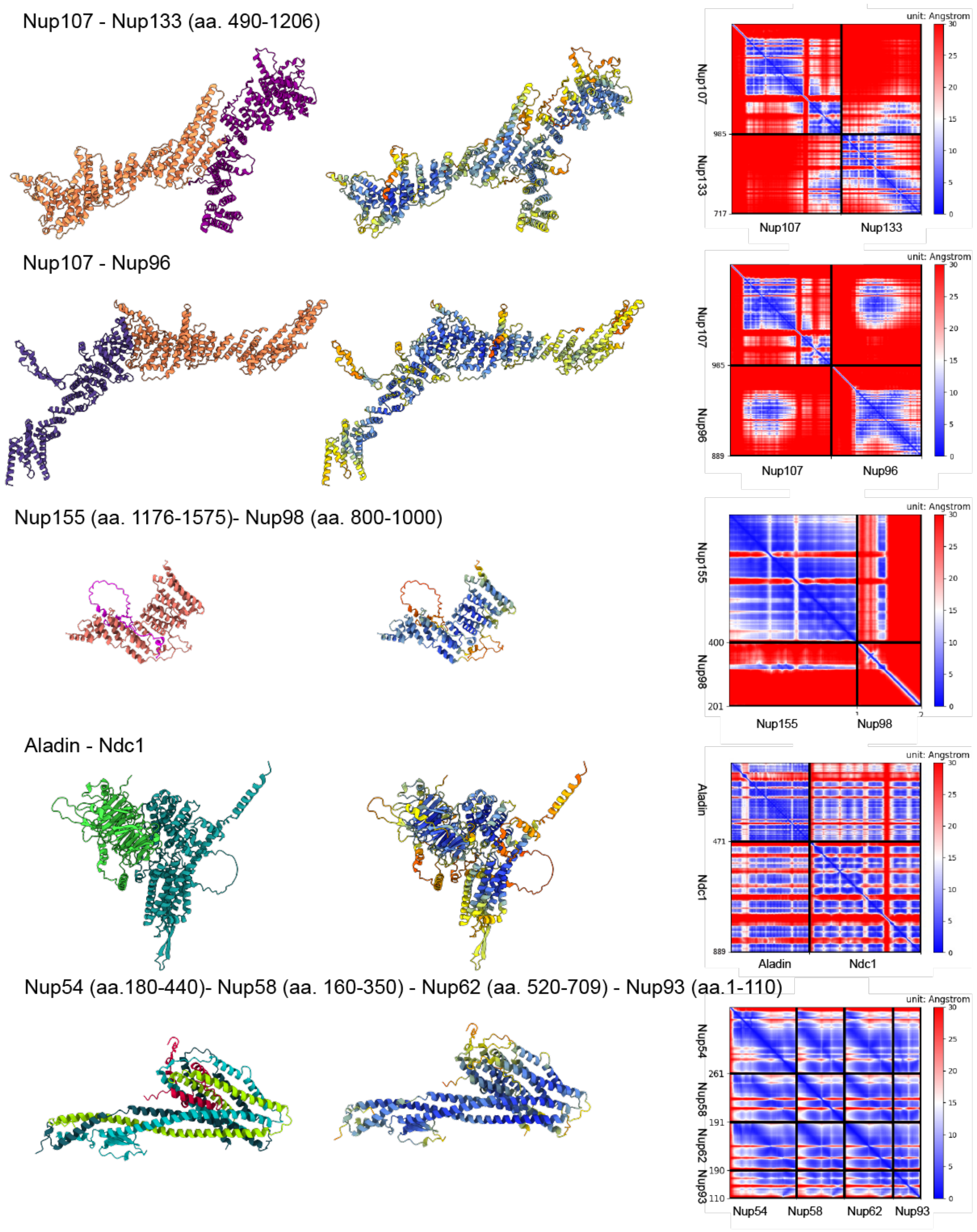
Nup107 - Nup133 (aa. 490-1206)
Nup107 - Nup96
Nup155 (aa. 1176-1575)- Nup98 (aa. 800-1000)
Aladin - Ndc1
Nup54 (aa.180-440)- Nup58 (aa. 160-350) - Nup62 (aa. 520-709) - Nup93 (aa.1-110)
unit: Angstrom
Nup107
Nup133
Nup96
Nup155
Nup98
Aladin
Ndc1
Nup54
Nup58
Nup62
Nup93

Nup35 (aa. 377-460) - Nup35 (aa. 377-460)

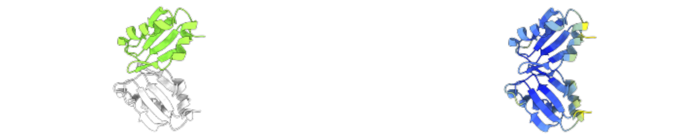

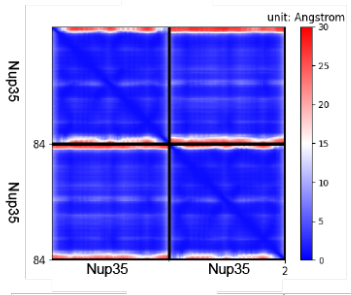


Nup93 (aa. 170-979) - Nup35 (aa.1-360)

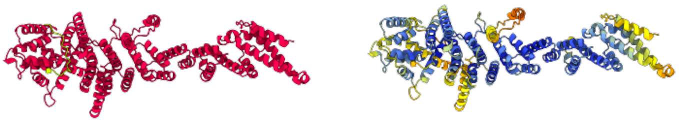

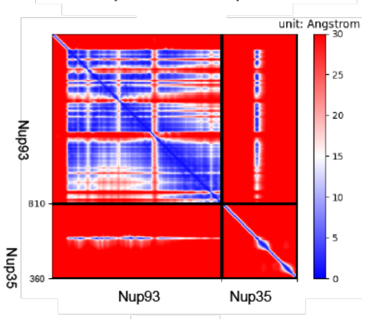


Nup205_N-Nup205_C - Nup93 (aa. 100-170)

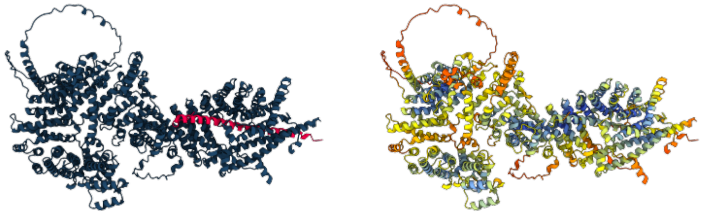

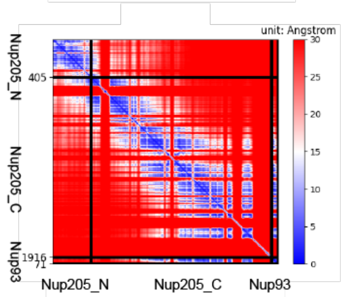


Nup188 - Nup93 (aa. 100-170)

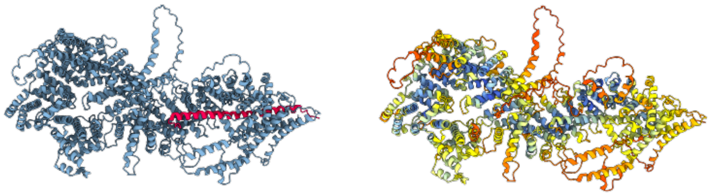

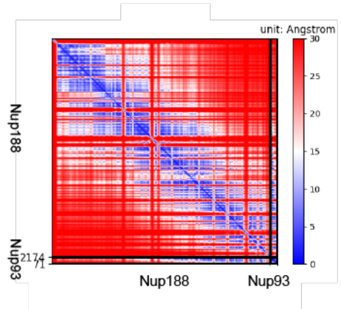


Nup214 (aa. 600-900) - Nup88 - Nup62 (aa. 521-709)

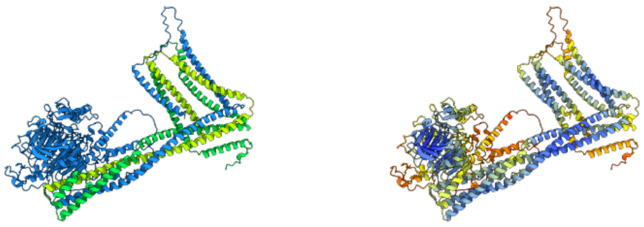

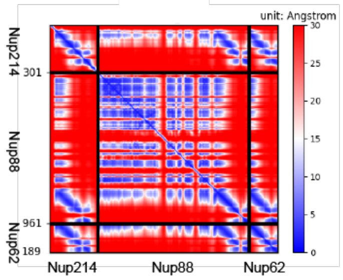

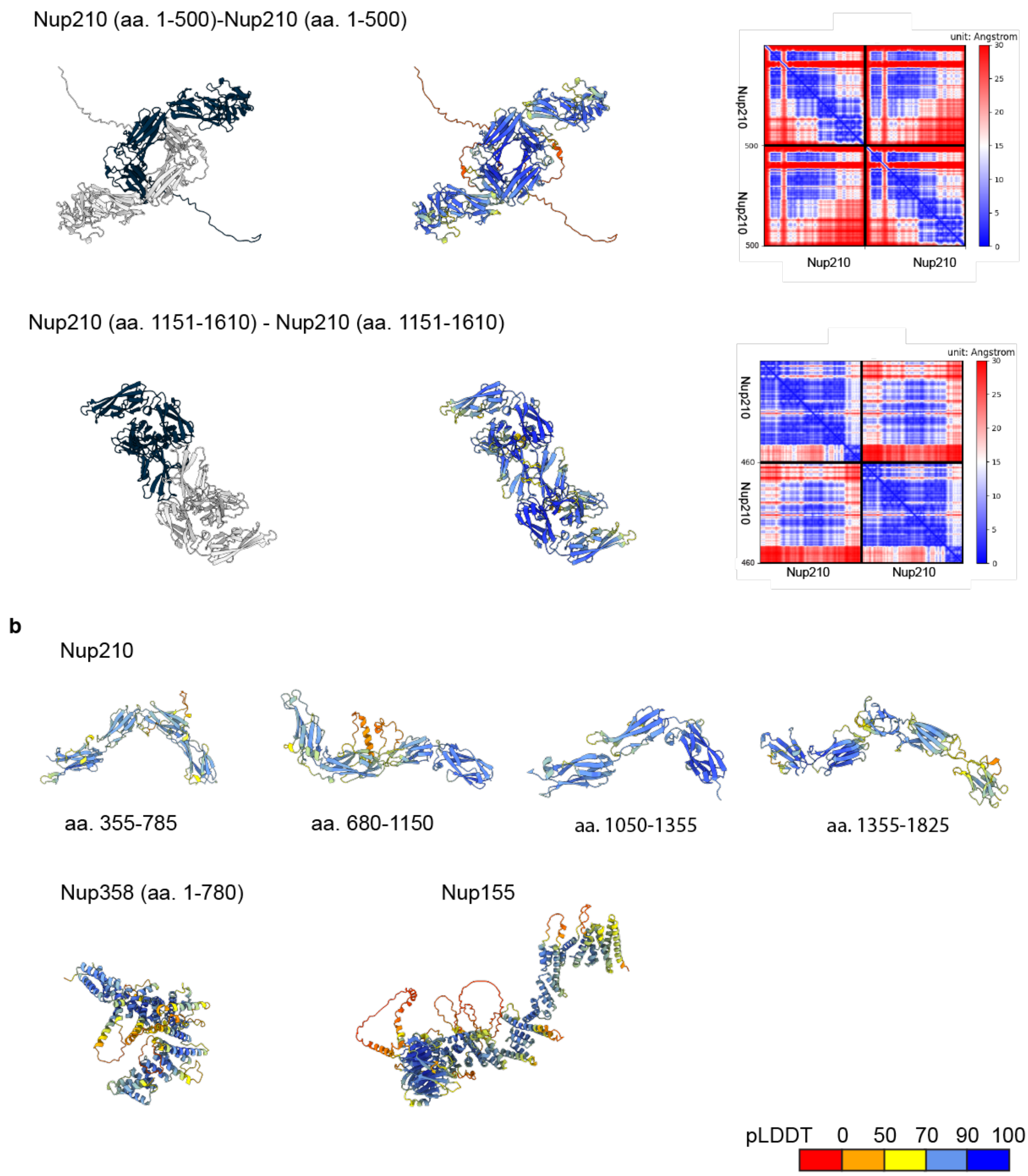


**Supplement Figure 5: Structural models of the *D. discoideum* Nups built using AlphaFold. a, Models of the Nup subcomplexes.** Each row shows, from left to right: a model colored by the Nup color code as shown in Fig. 3 and Fig. 4; a model colored by the local confidence estimated with predicted local distance difference test (pLDDT), as returned by AlphaFold. **b, Models of the monomeric Nups.** pLDDT > 90 (dark blue) indicates high estimated accuracy of backbone and side chain rotamers, whereas pLDDT > 70 (blue) indicates confident backbone prediction [59]; the confidence of interdomain and inter-chain orientations estimated with the expected distance error between all pairs of residues in the complex, as returned by AlphaPulldown/ColabFold. The color at each (x, y) position of the matrix corresponds to the expected distance error in residue x's position when the prediction and true (unknown) structure are aligned on residue y. Blue indicates low error. The heat maps show the sequence covered in the initial prediction and indicated in brackets (long disordered regions are not shown in the figures of model structures). aa. – amino acid residues.

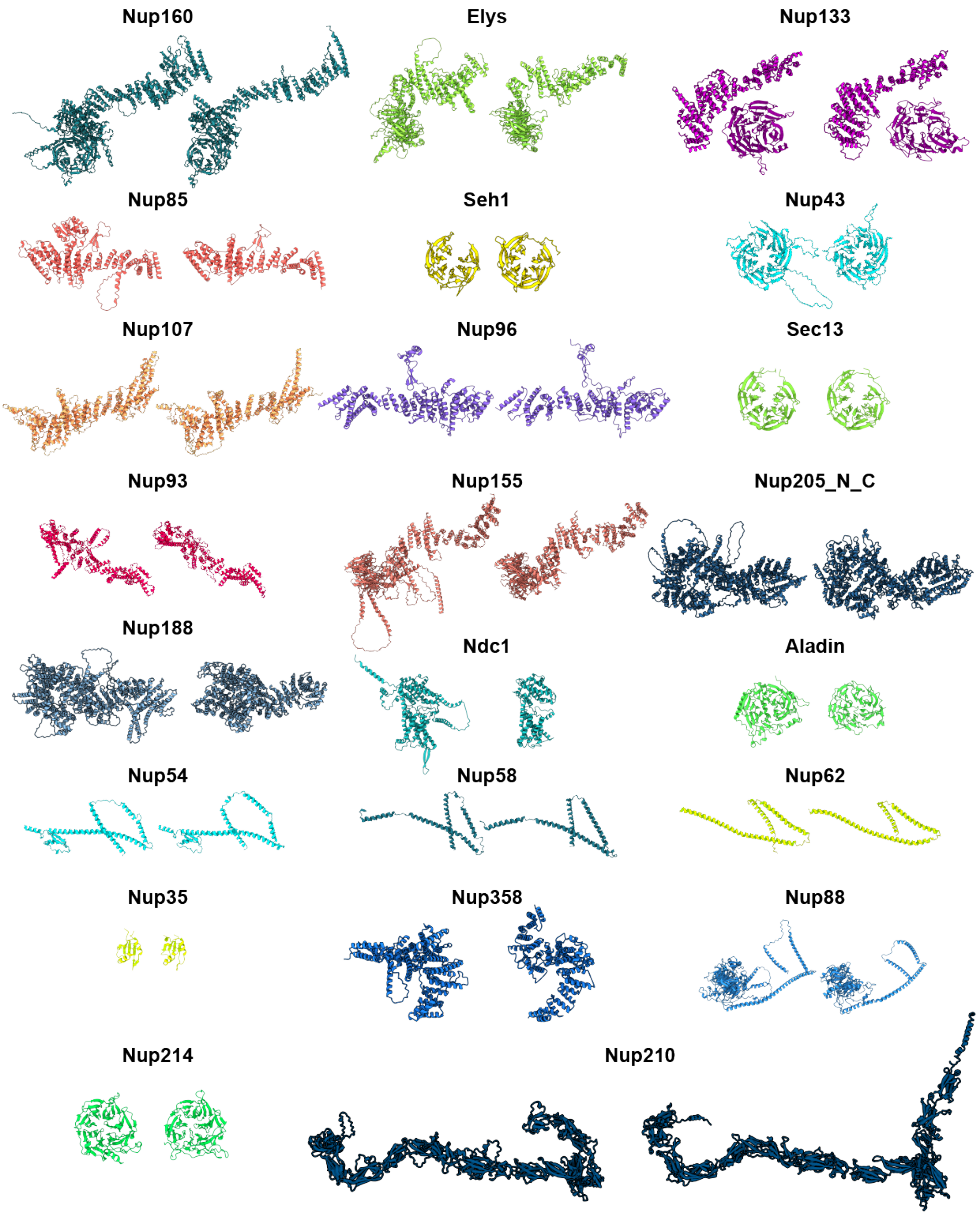


**Supplement Figure 6: Structural homologs of *D. discoideum* Nups detected by Foldseek.** On the left Nup from *D. discoideum*, on the right detected Nup listed in Supplement Table 4 Models are colored by the Nup color code as shown in Fig. 3 and Fig. 4.

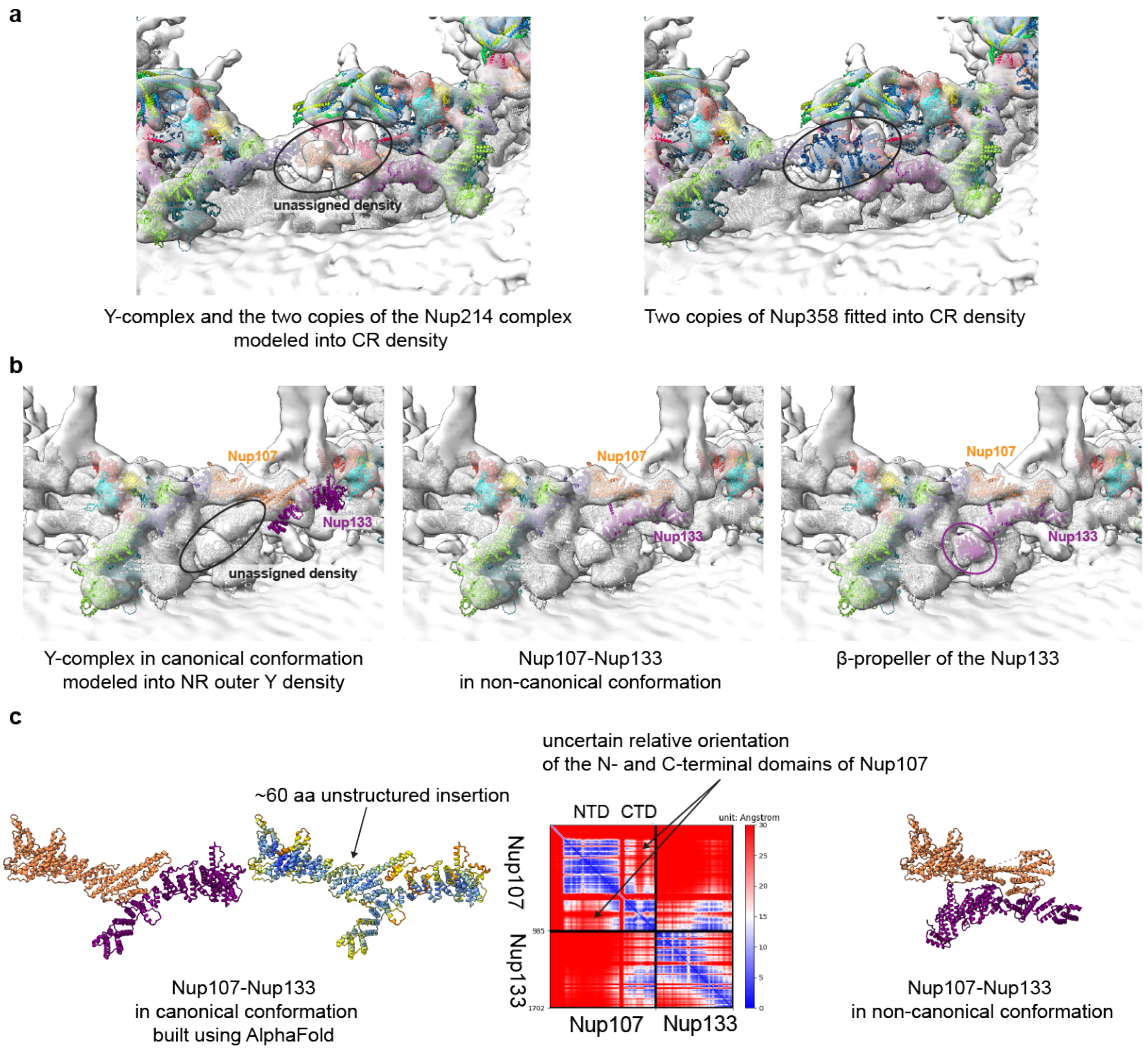


**Supplement Figure 7: Fitting of the *D. discoideum* Nups. a,** Fitted Y-complex and two copies of the Nup214 complex into CR revealed unassigned EM density *(left)*, into which we fitted two copies of the N-terminal domain of Nup358 *(right)*. **b,** *(left)* Modeling Y-complex in its canonical conformation in the third outermost Y-complex ring in the NR resulted in Nup133 placed outside of the EM density. However, the Nup133 structure could be fitted to the neighbouring unassigned density, suggesting alternative non-canonical conformation of the Nup107 (aa. 708-985)-Nup133 complex *(middle)*. *(right)* Resulting location of the Nup133 β-propeller. **c,** Nup107 (aa. 708-985)-Nup133 (aa. 490-1206) in its canonical and non-canonical conformation. Models are colored by the Nup color code as shown in Fig. 3 and Fig. 4 or by the pLDDT score, as in Supplement Fig. 5.

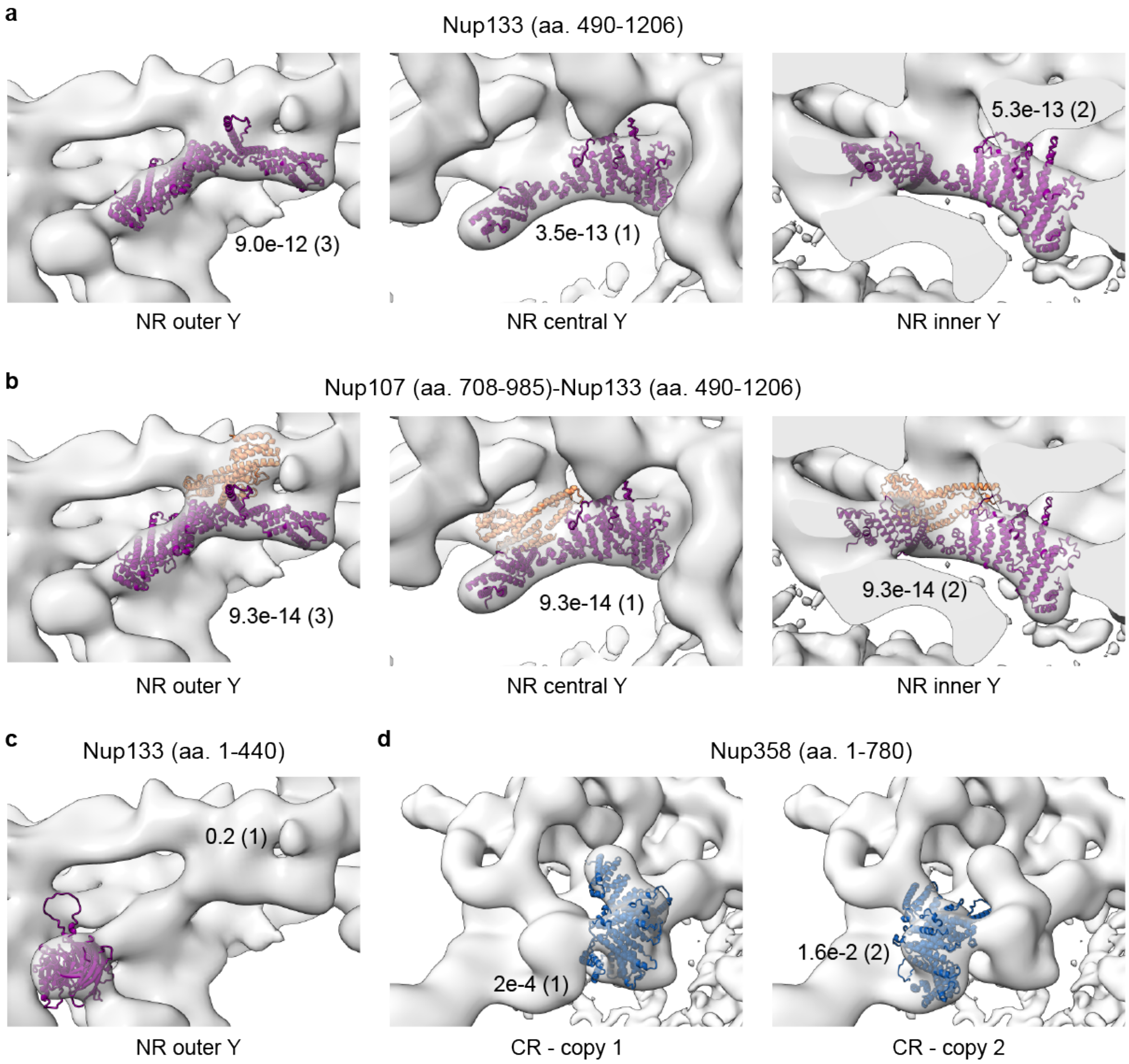


**Supplement Figure 8: Systematic fitting. a,** Nup133 (aa. 490-1206) fitted to the map of the NR assigned three copies of Nup133. **b,** Nup107 (aa. 708-985)-Nup133 (aa. 490-1206) fitted to the map of the NR assigned three copies of the Nup107-Nup133 subcomplex. **c,** β-propeller of the Nup133 fitted to the map of the NR outer. **d,** Nup358 fitted to the difference maps of the CR obtained after subtracting the density of Y-complex identifies two locations of Nup358. P-values, calculated as described in the Methods, and ranks of the fits are indicated next to the structures.

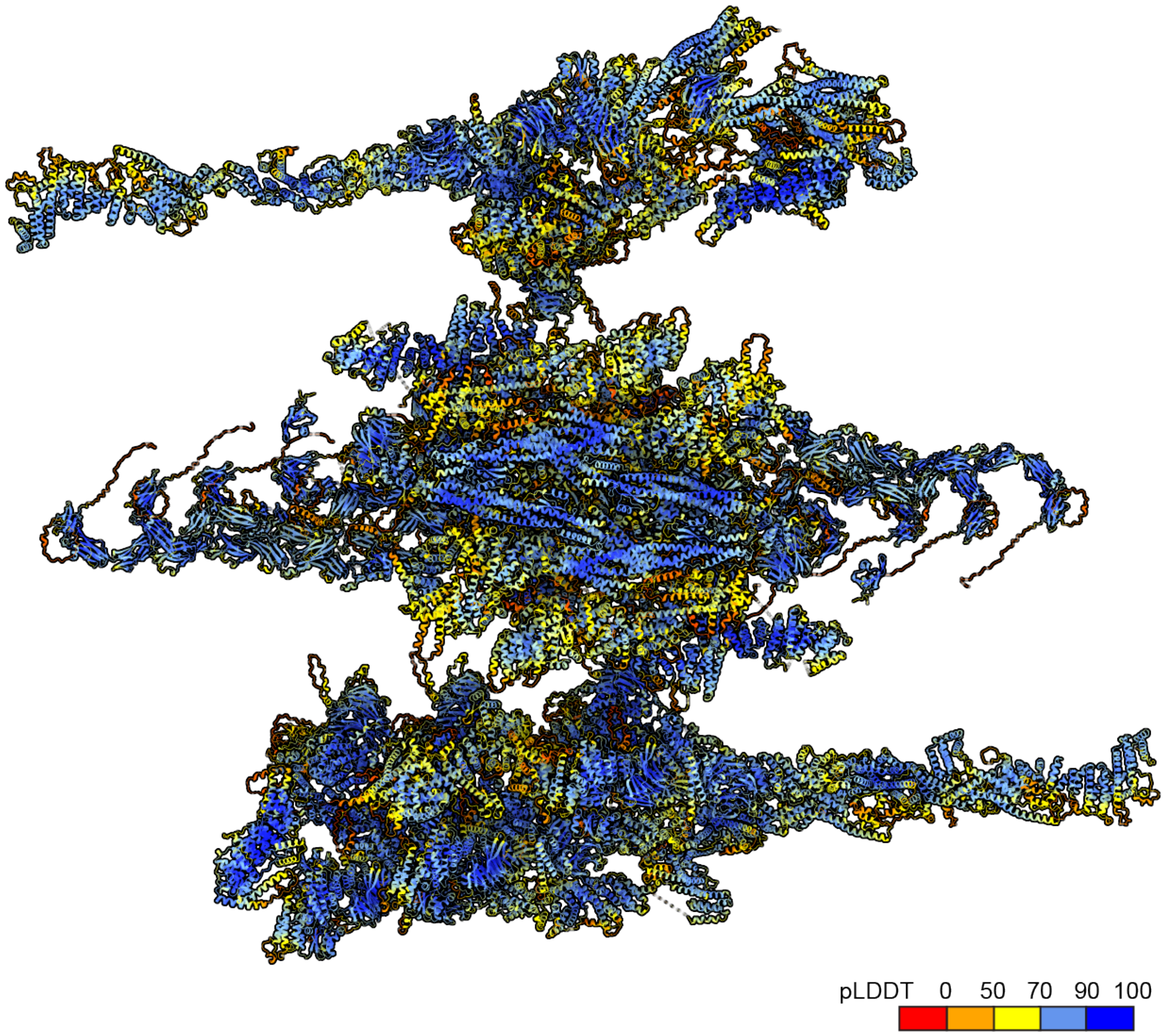


**Supplement Figure 9: Predicted local accuracy of the AlphaFold models projected onto an asymmetric unit of the NPC model.** The model is colored by local confidence estimated with predicted local distance difference test (pLDDT), as returned by AlphaFold. The pLDDT > 90 (dark blue) indicates high estimated accuracy of backbone and side chain rotamers whereas pLDDT > 70 (blue) indicates confident backbone prediction [59].

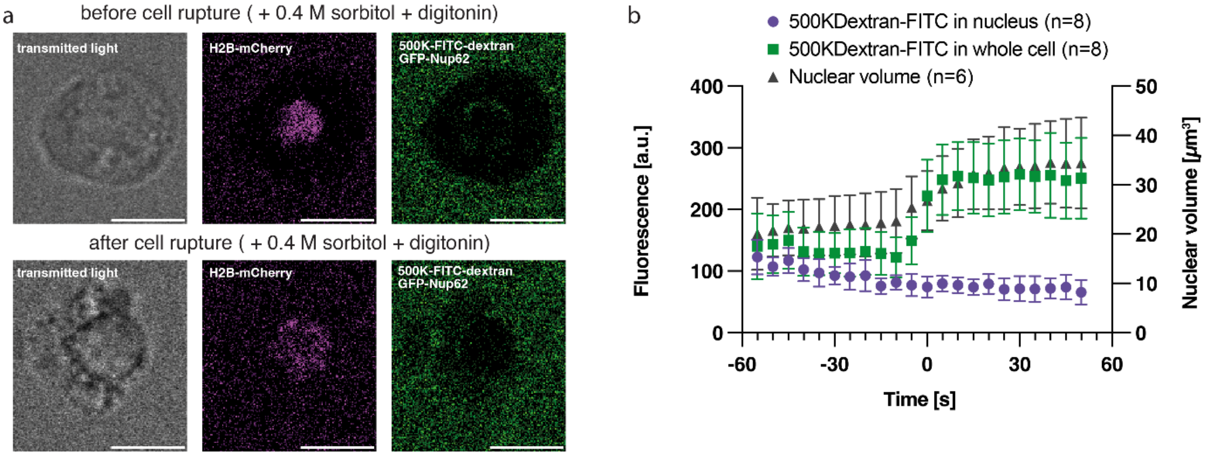


**Supplement Figure 10: The permeability barrier in *D. discoideum* during acute nuclear volume change is intact.**

**a,** Live cell images from confocal stacks of *D. discoideum* cell overexpressing the nuclear marker proteins GFP-Nup62 and mCherry-H2B in HL5 +0.4 M sorbitol + 20 µg/ml digitonin and 0.2 µM 500K-FITC-dextran. Cell is shown before and after plasma membrane rupture (respectively, top and bottom panels). Scale bar: 5 µm. **b,** Quantification of nuclear and cytosolic 500K-FITC-dextran signal during nuclear volume expansion when cells rupture in Hl5 + 0.4 M sorbitol + 20 µg/ml digitonin.

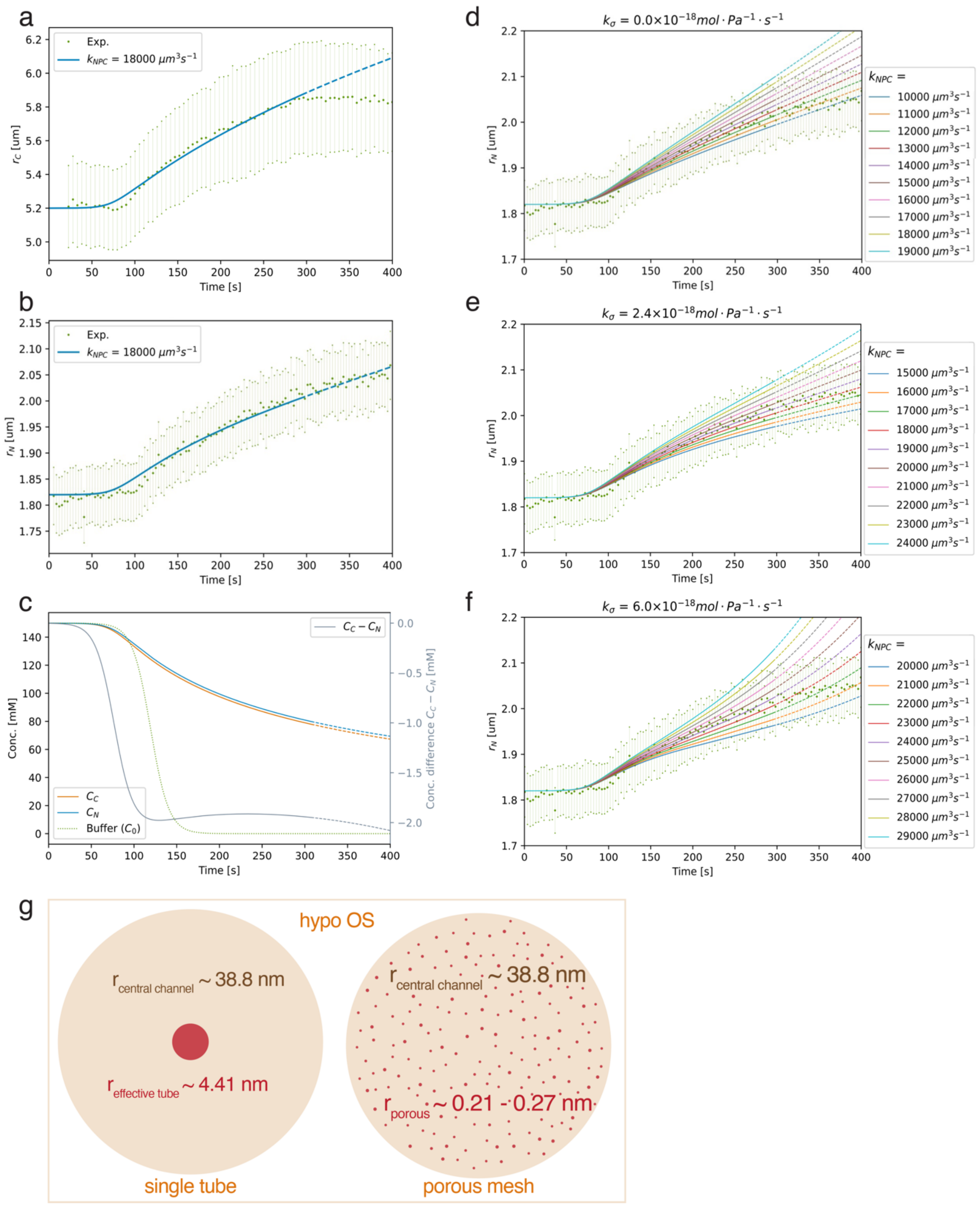


**Supplement Figure 11: Permeability of the NPC and corresponding uncertainty range during hypo OS.**

Experimental average and theoretically predicted time dependent **a,** cell and **b,** nucleus radii are plotted during hypo OS. Substituting the parameters described in Table S4 and $k_\sigma = 2.4 \times 10^{-18}\ mol \cdot Pa^{-1} \cdot s^{-1}$, in Eq. (10)-(13) in the Methods section, the permeability of one NPC $\left(k_{NPC_W}\right)$ was estimated to be 18000 $\mu m^3 \cdot s^{-1}$. **c,** Theoretically predicted time dependent concentrations of cytoplasm and nucleoplasm are plotted with experimentally determined buffer concentration for hypo OS. Shown on RHS vertical axis (gray) is the difference between the predicted concentrations. The optimum values for $k_\sigma$ and $k_{NPC_W}$ were determined by the Chi-Square test, within the range $k_\sigma \in [0 - 6 \times 10^{-18}]\, mol \cdot Pa^{-1} \cdot s^{-1}$ and $k_{NPC_W} \in [(1 - 3) \times 10^{-14}]\ m^3 \cdot s^{-1}$. Three example plots are shown for **d,** $k_\sigma = 0\, mol \cdot Pa^{-1} \cdot s^{-1}$, **e,** $k_\sigma = 2.4 \times 10^{-18}\, mol \cdot Pa^{-1} \cdot s^{-1}$ and **f,** $k_\sigma = 6 \times 10^{-18}\, mol \cdot Pa^{-1} \cdot s^{-1}$. **g,** Schematic display of central channel permeability from mathematical models during hypo OS based on experimental data. Left schematic shows the effective channel size radius of ~4.41 nm if one continuous single path through the FG mesh is assumed according to Hagen-Poiseuille law (model 1), model 2 shows the FG-mesh as continuous porous media with a critical radius of 0.21-0.27 nm according to Darcy's law.

**Supplement Table 1: Cryo-ET dataset properties.** Acquisition parameters and number of cells and tilt series used for NPC STA.

| | Control dataset | Hyper OS dataset | Hypo OS dataset |
|---|---|---|---|
| EMPIAR Deposition | EMPIAR-11845 | EMPIAR-XXX | EMPIAR-XXX |
| Microscope | Titan Krios G2 + FEG | Titan Krios G2 + FEG | Titan Krios G2 + FEG |
| Camera / EF | K3 / BioQuantum | K3 / BioQuantum | K3 / BioQuantum |
| Nominal Magnification | 42,000 | 42,000 | 42,000 |
| Pixel size (Å) | 2.176 | 2.176 | 2.176 |
| Defocus range (µm) | -2.5 to -5 | -2.5 to -5 | -2.5 to -5 |
| Dose rate (e/px/sec) | 10 - 20 | 10 - 20 | 10 - 20 |
| Total dose (e/Å$^2$) | ~132 - 150 | ~132 - 150 | ~132 - 150 |
| Number of cryo-ET sessions | 5 | 6 | 4 |
| Number of grids | 14 | 11 | 7 |
| Number milled lamellae on grids | 87 | 55 | 61 |
| Number of TS used for NPC processing | 133 of 247 total TS | 103 of 253 total TS | 74 of 163 total TS |
| Number of cells used for NPC processing | 47 | 52 | 24 |
| Number of NPCs | 348 | 248 | 149 |
| Number of NPC subunits | 2269 | 1683 | 969 |

**Supplement Table 2: Accession numbers of identified *D. discoideum* Nups by bioinformatics and included in the structural model.** The table displays the identified *D. discoideum* Nups in respect to their *H. sapiens* orthologs. The corresponding accession numbers and names were retrieved from the Uniprot database [149]. The presented Nups were modeled by AlphaFold and used to fit to the Cryo-ET maps.

| ***H. sapiens* Nup name** | ***D. discoideum* predicted Nup** | |
|---|---|---|
| | **UniProt ID** | **UniProt recommended name** |
| **Coat nucleoporin complex – Y-complex** | | |
| NUP160 | Q55DW5 | Uncharacterized protein |
| NUP85 | Q54NA0 | Nuclear pore complex protein nup85 |
| SEH1 | Q86K39 | Anaphase-promoting complex subunit 4-like WD40 domain-containing protein |
| NUP96 | Q54EQ8 | Nuclear pore complex protein Nup98-Nup96 |
| SEC13 | Q54DS8 | Protein SEC13 homolog |
| NUP107 | Q54N06 | Nuclear pore complex protein |
| NUP133 | Q54KA3 | Nucleoporin 133 |
| NUP43 | Q54Z22 | Nuclear pore complex protein nup43 |
| ELYS | Q55DZ3 | ELYS-like domain-containing protein |
| **Inner ring Nups** | | |
| NUP205_N | Q556V5 | Uncharacterized protein |
| NUP205_C | Q556V6 | Uncharacterized protein |
| NUP188 | Q54HL2 | Uncharacterized protein |
| NUP93 | Q55GW5 | Nuclear pore protein |
| NUP155 | Q54F20 | Nucleoporin 155 |
| NUP35 | Q75JH1 | RRM Nup35-type domain-containing protein |
| NUP54 | Q55BX5 | Nuclear pore complex protein nup54 |
| NUP58 | Q54XC5 | Uncharacterized protein |
| NUP62 | Q86IX4 | Nuclear pore protein |
| **POMs** | | |
| NDC1 | Q550C4 | Uncharacterized protein |
| NUP210 | Q54IS9 | Nucleoporin 210 |
| ALADIN | Q7KWY3 | WD40 repeat-containing protein |
| **Cytoplasmic filament Nups** | | |
| NUP88 | Q54XC9 | Uncharacterized protein |
| NUP214 | Q54H52 | WD40-like domain-containing protein |
| NUP358 | C7FZW3 | RanBP2-type zinc finger |

**Supplement Table 3: Physical parameters of protein-protein interfaces modelled by AlphaFold.**

The columns indicate as follows: “AlphaFold model”, “Interface” – the evaluated interface from the model, “Interface residues” – number of interface residues, “Polar”, ”Hydrophobic”, and “Charged” – fractions of the corresponding type of residues at the interface, “Contact pairs” – number of contact pairs, “Shape complementarity” – Shape complementarity, “Hydrogen bonds”, “Salt bridges” – number of potential salt bridges, “Solvation energy” – solvation free energy gain upon formation of the interface, “Interface area”, “P-value” – the P-value of the observed solvation free energy gain (P<0.5) implies higher likelihood that the interaction is specific [150]. Units are specified where applicable, “PI score” – PI score of the interface, with values > 0 indicating that the interface resembles real interfaces. All values were calculated with PI_score pipeline [151] using CCP4 [152], Rate4Site [153], and PISA [150] programs included in the pipeline. The models and were minimized using GROMACS prior to analysis.

| AlphaFold model | Interface | Interface residues | Polar | Hydro-phobic | Charged | Contact pairs | Shape compl. | Hydrogen bonds | Salt bridges | Solvation free energy [kcal/M] | Interface area [Å2] | P-value | PI score | TM score |
|---|---|---|---|---|---|---|---|---|---|---|---|---|---|---|
| **Nup107-Nup133 (490-1206)** | Nup107-Nup133 | 36 | 0.333 | 0.444 | 0.194 | 30 | 0.69 | 23 | 1 | -14.95 | 1648.49 | 0.26 | 2.21 | 0.65 |
| **Nup107-Nup96** | Nup107-Nup96 | 50 | 0.36 | 0.34 | 0.2 | 51 | 0.68 | 36 | 22 | -11.29 | 2235.71 | 0.54 | 1.54 | 0.60 |
| **Nup160-Elys (590-1170)-Nup96** | Nup160-Nup96 | 15 | 0.333 | 0.2 | 0.467 | 12 | 0.73 | 9 | 10 | -0.71 | 811.9 | 0.8 | 1.69 | 0.77 |
| **Nup160 (1-1200)-Elys (1-1100)** | Nup160-Elys | 35 | 0.457 | 0.171 | 0.286 | 27 | 0.53 | 22 | 12 | -9.52 | 2257.08 | 0.64 | 1.14 | 0.58 |
| **Nup160-Elys (590-1170)-Nup96** | Nup96-Elys | 18 | 0.667 | 0.278 | 0.056 | 13 | 0.74 | 15 | 2 | -12.53 | 1065.32 | 0.28 | 2.09 | *0.77* |
| **Nup160-Elys (590-1170)-Nup96** | Nup160-Elys | 58 | 0.414 | 0.241 | 0.259 | 50 | 0.57 | 31 | 8 | -13.74 | 2814.99 | 0.61 | 1.05 | 0.77 |
| **Nup160 (1120-1791)-Nup96-Sec13** | Nup96-Sec13 | 89 | 0.404 | 0.315 | 0.157 | 96 | 0.65 | 47 | 14 | -39.76 | 4172.66 | 0.15 | 1.02 | 0.72 |
| **Nup160 (1120-1791)-Nup96-Sec13** | Nup160-Nup96 | 65 | 0.415 | 0.262 | 0.277 | 59 | 0.64 | 46 | 24 | -18.02 | 3126.57 | 0.52 | 1.31 | 0.72 |
| **Nup160 (1000-1791)-Nup85-Seh1-Nup43** | Nup85-Nup43 | 26 | 0.5 | 0.231 | 0.192 | 21 | 0.61 | 31 | 15 | -1.36 | 1929.14 | 0.8 | 1.71 | 0.64 |
| **Nup160 (1000-1791)-Nup85-Seh1-Nup43** | Nup85-Seh1 | 28 | 0.357 | 0.321 | 0.143 | 21 | 0.67 | 12 | 13 | -15.5 | 1795.8 | 0.37 | 1.79 | 0.64 |
| **Nup160 (1000-1791)-Nup85-Seh1-Nup43** | Nup160-Nup85 | 39 | 0.436 | 0.308 | 0.179 | 26 | 0.74 | 30 | 13 | -11.0 | 1907.51 | 0.48 | 1.81 | 0.64 |
| **Nup214 (600-900)-Nup88-Nup62** | Nup214-Nup62 | 49 | 0.347 | 0.429 | 0.204 | 32 | 0.7 | 18 | 7 | -66.17 | 3610.7 | 0.03 | 1.2 | 0.58 |
| **Nup214 (600-900)-Nup88-Nup62** | Nup88-Nup62 | 76 | 0.237 | 0.5 | 0.211 | 54 | 0.69 | 47 | 23 | -59.73 | 4552.14 | 0.01 | 1.08 | 0.58 |
| **Nup214 (600-900)-Nup88-Nup62** | Nup214-Nup88 | 60 | 0.4 | 0.367 | 0.2 | 47 | 0.68 | 39 | 26 | -54.14 | 4213.44 | 0.02 | 1.06 | 0.58 |
| **Nup54(180-440)-Nup58(160-350)-Nup62(520-709)-Nup93(1-110)** | Nup54-Nup58 | 35 | 0.114 | 0.629 | 0.2 | 22 | 0.6 | 10 | 5 | -71.31 | 3712 | 0.01 | 1 | 0.79 |
| **Nup54(180-440)-Nup58(160-350)-Nup62(520-709)-Nup93(1-110)** | Nup58-Nup93 | 36 | 0.556 | 0.25 | 0.139 | 29 | 0.55 | 11 | 1 | -26.03 | 1907.2 | 0.47 | 1.04 | 0.79 |
| **Nup54(180-440)-Nup58(160-350)-Nup62(520-709)-Nup93(1-110)** | Nup54-Nup93 | 34 | 0.324 | 0.412 | 0.206 | 33 | 0.66 | 10 | 4 | -22.41 | 1442.49 | 0.33 | 2.55 | 0.79 |
| **Nup54(180-440)-Nup58(160-350)-Nup62(520-709)-Nup93(1-110)** | Nup62-Nup93 | 28 | 0.429 | 0.321 | 0.214 | 22 | 0.64 | 10 | 2 | -12.65 | 1242.4 | 0.65 | 1.88 | 0.79 |
| **Nup54(180-440)-Nup58(160-350)-Nup62(520-709)-Nup93(1-110)** | Nup54-Nup62 | 61 | 0.246 | 0.541 | 0.197 | 37 | 0.64 | 16 | 12 | -76.56 | 4195.74 | 0.07 | 1.14 | 0.79 |
| **Nup54(180-440)-Nup58(160-350)-Nup62(520-709)-Nup93(1-110)** | Nup58-Nup62 | 49 | 0.245 | 0.51 | 0.224 | 41 | 0.65 | 15 | 5 | -65.01 | 3475.38 | 0.03 | 1.24 | 0.79 |
| **Nup205-Nup93 (100-170)** | Nup205-Nup93 | 29 | 0.552 | 0.31 | 0.069 | 19 | 0.67 | 27 | 21 | -17.84 | 2491.28 | 0.73 | 1.03 | 0.81 |
| **Nup188-Nup93 (100-170)** | Nup188-Nup93 | 14 | 0.143 | 0.571 | 0.286 | 8 | 0.65 | 26 | 32 | -23.15 | 2797.26 | 0.64 | 1.02 | 0.86 |
| **Aladin-Ndc1** | Aladin-Ndc1 | 40 | 0.375 | 0.400 | 0.15 | 36 | 0.66 | 21 | 9 | -20.99 | 1925.21 | 0.27 | 1.82 | 0.83 |
| **Nup35(377-460)-Nup35(377-460)** | Nup35-Nup35 | 26 | 0.231 | 0.577 | 0 | 27 | 0.77 | 59 | 0 | 6.16 | 2153.85 | 0.97 | 0.92 | 0.87 |
| **Nup93 (170-979)-Nup35 (1-360)** | Nup93-Nup35 | 24 | 0.542 | 0.333 | 0.083 | 25 | 0.71 | 17 | 2 | -6.52 | 1077.01 | 0.08 | 2.32 | 0.60 |
| **Nup155 (1176-1575)-Nup98(815-915)** | Nup155-Nup98 | 30 | 0.4 | 0.4 | 0.133 | 27 | 0.65 | 22 | 11 | -21.81 | 2172.99 | 0.07 | 1.6 | 0.67 |
| **Nup210 (1-500)-Nup210 (1-500)** | Nup210-Nup210 | 44 | 0.409 | 0.318 | 0.227 | 48 | 0.69 | 34 | 10 | -2.34 | 1655.87 | 0.94 | 1.95 | 0.69 |
| **Nup210 (1151-1610)-Nup210 (1151-1610)** | Nup210-Nup210 | 46 | 0.348 | 0.304 | 0.174 | 47 | 0.69 | 23 | 10 | -24.0 | 1969.07 | 0.12 | 2.08 | 0.77 |

**Supplement Table 4: Foldseek results.** The columns indicate as follows: Model of *D. discoideum* Nup used as a query, name of the target structure detected by Foldseek, PDB code containing the target structure, organism, probability, sequence identity, TM-Score[1] normalized by the query length, TM-Score[2] normalized by the average length of query and target, RMSD between query and the target, residue ranges of the query and the target used to calculate TM-Scores and RMSD. "-" means that Foldseek didn't find any target, or TM-score didn't work, "*" means that TM-score[2] and RMSD were calculated using TM-score server (https://zhanggroup.org/TM-score/ [154,155]), when Foldseek failed to provide these values.

| ***D.discoideum* Nup – query** | **Target name** | **PDB code** | **Organism** | **Prob.** | **Seq. Id.** | **TM-score[1]** | **TM-Score[2]** | **RMSD** | **Query residues** | **Target residues** |
|---|---|---|---|---|---|---|---|---|---|---|
| Nup160 | Nup160 | 7WB4 | *H.sapiens* | 0.82 | 11.2 | 0.727 | *0.78 | *4.71 | 23-1791 | 1-1297 |
| Elys | Elys | 7R5K | *H.sapiens* | 0.82 | 10.5 | 0.656 | 0.72 | 6.01 | 6-1100 | 5-943 |
| Nup133 | Nup133 | 7TDZ | *X.laevis* | 0.89 | 11.8 | 0.681 | 0.70 | 4.72 | 490-1206 | 1-671 |
| Nup133 | Nup133 | 1XKS | *H.sapiens* | 0.95 | 12.8 | 0.755 | 0.79 | 3.22 | 1-404 | 5-374 |
| Nup85 | Nup85 | 7FIK | *X.laevis* | 0.72 | 15.5 | 0.712 | 0.85 | 3.57 | 90-890 | 1-593 |
| She1 | She1 | 6LK8 | *X.laevis* | 1.00 | 53.5 | 0.913 | 0.93 | 1.41 | 1-204,365-467 | 1-293 |
| Nup43 | Nup43 | 7R5J | *H.sapiens* | 0.93 | 16.1 | 0.779 | *0.71 | *2.57 | 2-417 | 1-343 |
| Nup107 | Nup107 | 7WB4 | *X.laevis* | 0.99 | 23.4 | 0.833 | 0.85 | 3.52 | 126-650,694-985 | 1-785 |
| Nup96 | Nup96 | 7WB4 | *X.laevis* | 0.99 | 19 | 0.828 | 0.84 | 3.48 | 211-889 | 1-672 |
| Sec13 | Sec13 | 7TDZ | *X.laevis* | 1.00 | 55.9 | 0.943 | 0.94 | 1.83 | 1-301 | 3-305 |
| Nup93 | Nup93 | 7WB4 | *X.laevis* | 0.97 | 16.4 | 0.808 | 0.86 | 3.09 | 251-979 | 1-641 |
| Nup155 | Nup155 | 7TBI | *S. cerevisiae* | 0.89 | 12.8 | 0.747 | *0.65 | *5.21 | 161-1564 | 1-1231 |
| Nup205 | Nup205 | 7VCI | *X.laevis* | 0.87 | 9.6 | 0.699 | *0.78 | *5.58 | 1-2128 | 10-1923 |
| Nup188 | Nup188 | 7MVX | *C.thermophilum* | 0.66 | 5.3 | 0.649 | 0.73 | 6.54 | 38-2080 | 1-1614 |
| Ndc1 | Ndc1 | 7WKK | *X.laevis* | 0.69 | 9.7 | 0.712 | 0.85 | 3.5 | 41-660 | 1-431 |
| Aladin | Aladin | 7WKK | *X.laevis* | 0.92 | 19.1 | 0.789 | 0.88 | 2.64 | 5-470 | 1-367 |
| Nup54 | Nup54 | 5CWS | *C.thermophilum* | 0.95 | 18.1 | 0.722 | 0.72 | 3.88 | 199-440 | 1-241 |
| Nup58 | Nup58 | 5CWS | *C.thermophilum* | - | - | - | 0.65 | 3.90 | 170-349 | 1-180 |
| Nup62 | Nup62 | 5CWS | *C.thermophilum* | - | - | - | 0.77 | 2.73 | 529-709 | 1-241 |
| Nup35 | Nup35 | 1WWH | *M.musculus* | 0.99 | 30.8 | 0.830 | 0.84 | 1.92 | 3-81 | 1-79 |
| Nup358 | Nup358 | 7FIK | *X.laevis* | 0.28 | 7.1 | 0.470 | 0.59 | 6.34 | 15-706 | 10-556 |
| Nup88 | Nup88 | 7VOP | *X.laevis* | 0.41 | 7.4 | 0.552 | 0.62 | 5.93 | 1-961 | 1-718 |
| Nup214 | Nup214 | 3FMP | *H.sapiens* | 0.99 | 14.6 | 0.82 | 0.85 | 2.96 | 1-430 | 1-411 |
| Nup210 | Nup210 | 7R5J | *H.sapiens* | 0.51 | 9.9 | 0.527 | - | - | 46-1713 | 32-1678 |

**Supplement Table 5: Input variables and corresponding uncertainty ranges** used in Eq. (10)-(13), to predict the permeability of the NPCs. Experimentally determined uncertainty ranges are marked (Exp).

| **Predetermined Variables** | **Value Used** | **Predetermined Error Range** | **Explanation** |
|---|---|---|---|
| $r_{C_0}$ | $5.25\ \mu m$ | $4.92 - 6.20$ (Exp) | Initial cell radius (hyper OS) |
| $r_{C_0}$ | $5.1\ \mu m$ | $4.06 - 5.75$ (Exp) | Initial cell radius (hypo OS) |
| $r_{N_0} = R_N$ | $1.58\ \mu m$ | $1.44 - 1.75$ (Exp) | Initial nucleus radius (hyper OS) |
| $r_{N_0} = R_N$ | $1.82\ \mu m$ | $1.60 - 1.97$ (Exp) | Initial nucleus radius (hypo OS) |
| $C_{C_0}$ | $150\ mole/m^3$ | Negligible | Initial cyto conc. |
| $C_{N_0}$ | $150\ mole/m^3$ | Negligible | Initial nucleus conc. |
| $C_O(t)$ | Exp. Avg. | $\sim \pm 20\%$ (Exp) | Buffer conc. |
| $v_W$ | $1.8 \times 10^{-5}\ m^3$ $/mole$ | Negligible | Water vol. per mole |
| $k_{mem}$ | $3.5 \times 10^{-5}\ m/s$ | $3 - 4 \times 10^{-5}$ | Cell permeability |
| $k_{mem-NE}$ | $3.5 \times 10^{-4}\ m/s$ | $1.75 - 7 \times 10^{-4}$ | NE permeability |
| $V_{C-exc}$ | $0.5(V_{cell} - V_N)$ | $50 \pm 5\%$ | Excluded cyto vol. |
| $V_{N-exc}$ | $0.5V_N$ | $50 \pm 20\%$ | Excluded nucleus vol. |
| $N_{NPC}$ | 450 | 380-540 | Number of NPCs |
| $\sigma_{NE}$ | $0.05\text{Nm}^{-1}$ | $0.025\text{-}0.1\text{Nm}^{-1}$ | Nuclear envelope tension |
